# Missense mutations in CRX homeodomain cause dominant retinopathies through two distinct mechanisms

**DOI:** 10.1101/2023.02.01.526652

**Authors:** Yiqiao Zheng, Chi Sun, Xiaodong Zhang, Philip A. Ruzycki, Shiming Chen

**Author notes:** To whom the correspondence should be addressed. Mailing address: 660 South Euclid Avenue, MSC-8096-06-06, St. Louis, MO 63110, USA.

## Abstract

Homeodomain transcription factors (HD TFs) are instrumental to vertebrate development. Mutations in HD TFs have been linked to human diseases, but their pathogenic mechanisms remain elusive. Here we use *Cone-Rod Homeobox (CRX)* as a model to decipher the disease-causing mechanisms of two HD mutations, *p.E80A* and *p.K88N*, that produce severe dominant retinopathies. Through integrated analysis of molecular and functional evidence *in vitro* and in knock-in mouse models, we uncover two novel gain-of-function mechanisms: *p.E80A* increases CRX-mediated transactivation of canonical CRX target genes in developing photoreceptors; *p.K88N* alters CRX DNA-binding specificity resulting in binding at ectopic sites and severe perturbation of CRX target gene expression. Both mechanisms produce novel retinal morphological defects and hinder photoreceptor maturation distinct from loss-of-function models. This study reveals the distinct roles of E80 and K88 residues in CRX HD regulatory functions and emphasizes the importance of transcriptional precision in normal development.

## Introduction

Homeodomain transcription factors (HD TFs) play a fundamental role in vertebrate development. Members of the HD TF family are characterized by the presence of a highly conserved 60 amino acid helix-turn-helix DNA-binding domain known as the homeodomain (HD). The HD is one of the most studied eukaryotic DNA-binding motifs since its discovery in *Drosophila* homeotic transformations^1,2^. Hundreds of HD TFs have subsequently been documented in regulating gene expression programs important for body plan specification, pattern formation and cell fate determination^1,2^. Mutations in HD TFs have been linked to many human diseases, including neuropsychiatric and neurodegenerative conditions^3,4^. Although significant progress has been made in understanding HD-DNA interactions, uncovering the pathogenetic mechanisms of disease-causing missense mutations in HD have proven challenging.

The retina has long been used as a model system to study the role of HD TFs in normal central nervous system (CNS) development and in neurological diseases^5^. During retinogenesis, HD TFs play essential roles in the patterning of neuroepithelium, specification of retinal progenitors and differentiation of all retinal cell classes that derive from a common progenitor^6^. Importantly, many HD TFs are shared between the brain and the retina during development and mutations in these TFs can lead to disease manifestation in both tissues^7–12^. The accessibility and wealth of available molecular tools makes the retina a valuable tool to decipher the pathogenic mechanisms of HD TF mutations associated with neurological diseases.

Here, we study CRX, a HD TF essential for photoreceptor cells in the retina, as a model to understand how single amino acid substitutions in the HD impact TF functions and cause blinding diseases. Photoreceptors are the most numerous neurons in the retina and are specialized to sense light and initiate vision through a process called phototransduction. Animal studies have demonstrated that *Crx* is first expressed in post-mitotic photoreceptor precursors^13^ and maintained throughout life^14,15^. Loss of CRX results in impaired photoreceptor gene expression, failure of maturation and rapid degeneration of immature, non-functional photoreceptors^16^. Protein-coding sequence variants in human *CRX* have been associated with inherited retinal diseases (IRDs) that affect photoreceptors: Leber congenital amaurosis (LCA), cone-rod dystrophy (CoRD), and retinitis pigmentosa (RP) (OMIM:602225). However, the recessive phenotype observed in *Crx* knockout mouse models fails to recapitulate many dominant human *CRX* mutations that arise *de novo*^16^.

CRX contains two functional domains – the N-terminal HD and C-terminal activation domain (AD) (Figure 1A), both are required for proper activation of target genes and maintenance of normal *Crx* mRNA transcript abundance^17,18^. To understand how CRX HD mutations cause diseases, we have previously reported a mutation knock-in mouse model carrying a hypomorphic mutation *p.R90W* (R90W) in CRX HD^19,20^. We found that R90W mutation produces a recessive loss-of-function phenotype very similar to that of *Crx* knockout mice.

**Figure 1.**
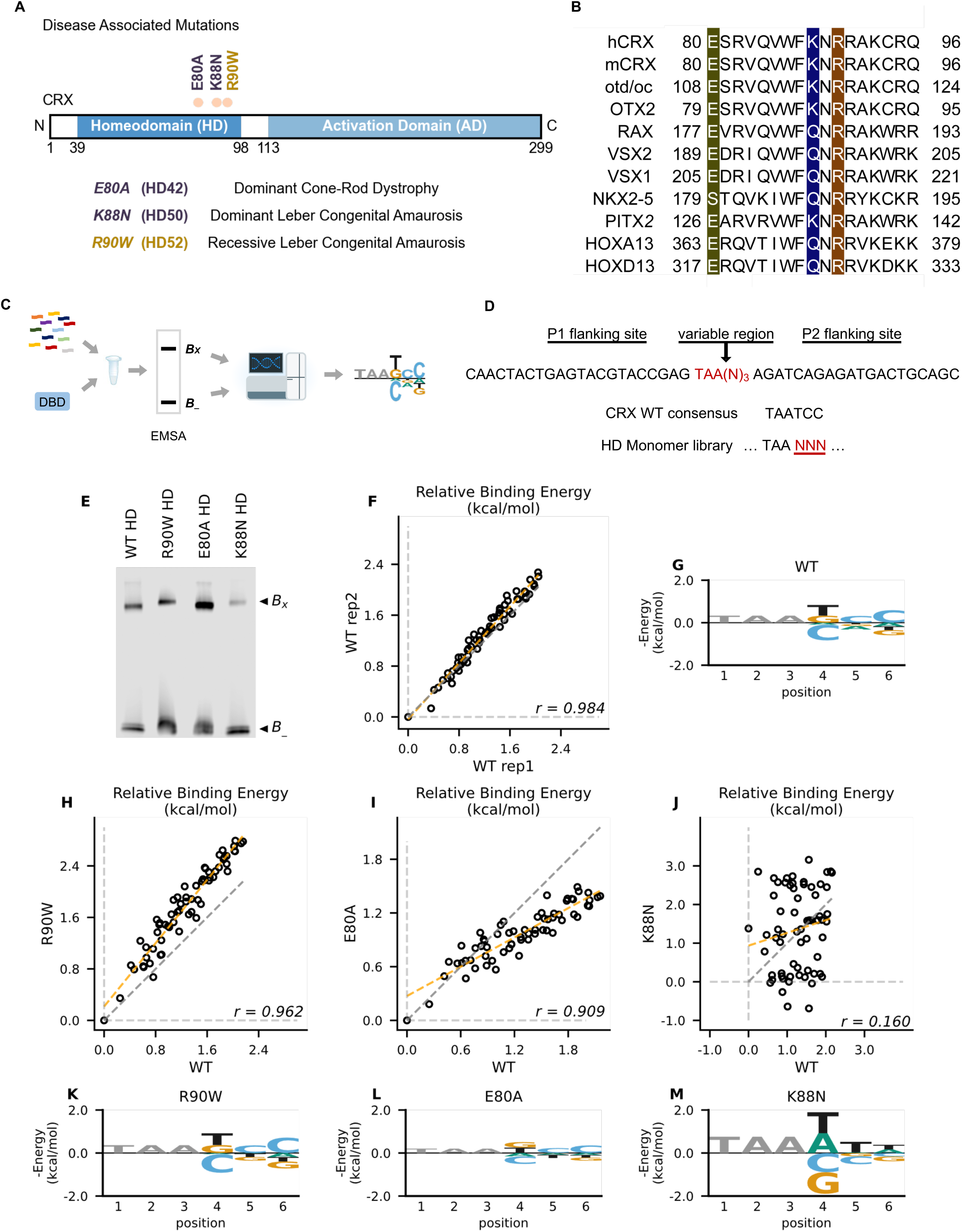
Disease associated missense mutations altered CRX HD DNA binding specificity. (A) Diagram of CRX functional domains: Homeodomain (HD) for DNA-binding and Activation Domain (AD) for target gene transactivation. The three missense mutations in this study are located at the C-terminus of CRX HD and associated with different retinal diseases in human. Number in the parenthesis denotes the CRX HD position of the corresponding mutated residue. (B) Alignments of HD recognition helix sequences for the indicated HD proteins for which HD missense mutations have been associated with inherited diseases. Accession numbers can be found in Supplementary Table S1. Missense variants in this study (highlighted) are located at highly conserved residues across species and different HD TFs. (C) Spec-seq experimental workflow (Methods). (D) Spec-seq library design of monomeric HD binding sites. (E) EMSA gel images of Spec-seq experiments with different CRX HD species. Bx: Bound. B-: Unbound. (F) Relative binding energy comparison from two different experiments with WT HD. (G) Binding energy model for WT CRX HD. (H-J) Relative binding energy comparison between WT HD and R90W HD (H), E80A HD (I), or K88N HD (J). Consensus sequence is defined to have relative binding energy of 0kT (TAATCC for WT, R90W and E80A, TAATTA for K88N). The identity line is represented in grey dash. The orange dashed line shows the best linear fit to the data. (K-M) Binding energy models for R90W HD (K), E80A HD (L), and K88N HD (M). Only sequence variants within two mismatches to the corresponding consensus sequences were used to generate binding models. Negative binding energy is plotted such that bases above the x-axis are preferred bases and bases below the x-axis are unfavorable bases. Constant bases (TAA) carried no information are drawn at arbitrary height in grey.

Intriguingly, several missense mutations within the same HD recognition helix as R90W, including *p.E80A* (E80A) and *p.K88N* (K88N), are linked to severe dominant IRDs^21–23^ (Figures 1A and 1B). Here, we utilized a multi-omics approach to investigate the functional consequences of the E80A and K88N mutations on CRX regulatory activities and photoreceptor development (Figure S1). Comparison of the *in vitro* HD-DNA binding models of CRX and disease variants generated by Spec-seq revealed unique specificity changes of each mutant protein. Introduction of each mutation into the endogenous *Crx* locus generated knock-in mouse models *Crx^E80A^* and *Crx^K88N^* that reproduced dCoRD- and dLCA-like phenotypes. ChIP-seq analysis of CRX binding *in vivo* revealed mutation-specific changes in CRX targetome, consistent with mutation-specific DNA binding changes *in vitro*. Retinal RNA-seq analysis uncovered two distinct mechanisms by which the two HD missense mutations contribute to altered gene expression programs during photoreceptor differentiation and maturation. Our results highlight the importance of residues E80 and K88 in CRX-mediated transcriptional regulation during photoreceptor development and the diverse mechanisms by which HD missense mutations can affect TF functions and lead to severe dominant neurological diseases.

## Results

### K88N but not E80A mutation alters CRX HD DNA-binding specificity *in vitro*

CRX belongs to the *paired*-like HD TF family that recognize a 6-bp DNA motif in a stereotypic way^24–30^. Extensive studies of the HD have revealed a canonical HD-DNA recognition model where recognition of the 3’ region (bases 4-6) of the HD DNA binding site is mediated by specificity determinants within the conserved HD recognition helix, corresponding to CRX residues 80-96^24–27,30,31^ (Figure 1B). In particular, HD residue 50, equivalent to CRX K88 residue (Figure 1A), is the major specificity determinant in *paired*-like HD TF-DNA interactions^25,26^. Since the three disease-associated HD missense mutations, E80A, K88N and R90W, are located within the CRX HD recognition helix, we wondered if these mutations change CRX HD DNA-binding specificity.

We adapted a high-throughput *in vitro* assay, Spec-seq, that determines protein-DNA-binding specificity by sequencing^32–34^. Spec-seq was developed based on the traditional electrophoretic mobility shift assay (EMSA) to measure protein-DNA interactions. Spec-seq allows us to measure the relative binding affinities (i.e., specificity) for a library of HD binding motifs in parallel and generate quantitative binding models for different CRX HDs (Figures 1C). Based on the HD-DNA interaction model, we designed and tested a Spec-seq library containing all possible monomeric HD motifs (TAANNN) (Figure 1D).

We first obtained the wild-type (WT) CRX HD DNA binding model with Spec-seq using bacterially-expressed and affinity-purified HD peptides (Figures 1E-1G, Methods). Relative binding energies of CRX WT HD from two experiments showed strong correlation (*r*: 0.984) and noise level (0.114 kT) within the expected range in typical Spec-seq data (Figure 1F). Binding energy model of WT HD was then generated by applying multiple linear regressions on the relative binding energies of all sequences within two base-pair mismatches to the WT CRX consensus (TAATCC)^14^ (Figure 1G, Methods). A clear preference for CC bases at the 3’ end of the motif is consistent with known CRX binding preference *in vitro* and *in vivo*^35,36^.

We next sought to understand how disease mutations affect CRX DNA-binding specificity. We purified all mutant HD peptides following the same protocol as WT HD peptides and verified their DNA binding (Figures S2A-S2D)^18^. Comparison of the relative binding energies between each pair of mutant and WT HD revealed distinct effects (Figures 1H-1J). By definition, the consensus DNA binding motif of a testing peptide has a relative binding energy of 0kT. The relative binding energy difference between nucleotide variants and the consensus motif correlates with the DNA binding specificity of the testing peptide. We found that R90W HD and E80A HD both prefer the same consensus motif as WT (Figures 1K and 1L). R90W HD bound with slightly higher specificity than WT, as demonstrated by most data points falling above the identity line (Figures 1H and 1K), suggesting that R90W HD is more sensitive than WT to binding sequence variations. In contrast, when comparing E80A HD with WT HD, many data points fell below the identity line and the relative binding energies regressed towards 0 on the E80A axis (Figures 1I and 1L). This suggests that E80A HD bound with lower specificity than WT HD and thus was more tolerant to base variations in the HD DNA motif. Different from R90W and E80A, K88N mutation dramatically altered CRX HD DNA-binding specificity (*r*: 0.160) (Figures 1J and 1M). The K88N preferred binding sequence (TAAT/ATT/A) is referred to as N88 HD motif hereafter. K88N HD also had the largest degree of discrimination from its preferred to the weakest binding motif, suggesting that it is most sensitive to variants in the HD DNA motif. As a control, we tested a second library with the TAANNN sites on the reverse strand and obtained similar results (Figures S2E-S2L). Together, these results indicate that while E80A mutation does not affect CRX HD DNA binding specificity, the K88N mutation dramatically alters the specificity *in vitro*.

### E80A protein binds to WT sites while K88N occupies novel genomic regions with N88 HD motifs *in vivo*

Next, we asked if changes in DNA-binding specificity affected mutant CRX chromatin binding in developing photoreceptors. We first created two human mutation knock-in mouse models, *Crx^E80A^* and *Crx^K88N^*, each carrying a single base substitution at the endogenous *Crx* locus (Figures S3A and S3B, Methods). We confirmed that *Crx* mRNA was expressed at comparable levels in WT and mutant retinas (Figure S3C), and the full-length CRX proteins were readily detectable in the nuclear extracts from all samples (Figure S3D). We then obtained genome-wide binding profiles for each CRX variant by chromatin immunoprecipitation followed by sequencing (ChIP-seq) on mouse retinas at P14, a time when all retinal cell types are born, photoreceptor specification is completed in *WT* animals, and prior to any observed cell death in other CRX mutants previously characterized^19,37^. To focus on changes specific to each mutant CRX protein, only homozygous animals were used for ChIP-seq profiling.

Unsupervised clustering of all CRX binding sites revealed two major clusters (Figure 2A, Methods). Cluster 1 consisted of canonical WT CRX binding sites that are also occupied by CRX E80A protein (Figures 2A and 2B). Similar to WT CRX, CRX E80A binding *in vivo* was mostly enriched in intronic, followed by intergenic and TSS regions (Figure 3D). In contrast, CRX R90W, that also showed similar consensus preference to WT *in vitro*, failed to produce significant DNA binding *in vivo* (Figures 2A-2C, Methods). This suggests that the retinopathy phenotype of *Crx^R90W/W^* is likely due to loss of binding at canonical WT CRX binding sites. Intriguingly, while CRX K88N showed loss of binding at canonical CRX binding sites, it gained a small set of binding sites (Cluster 2, Figures 2A-2C). *De novo* motif searching with DREME^38^ under CRX peaks in each genotype revealed enrichment of monomeric HD motifs (Figure 2E) consistent with those found in Spec-seq (Figures 1G and 1K-1M), highlighted by a change in enriched HD motif from WT CRX HD type to N88 HD type in the *Crx^K88N/N^* retinas. Consistency with *in vitro* binding models suggests that *in vivo* changes in CRX chromatin binding were at least in part driven by the intrinsic changes in HD-DNA binding specificity by each individual mutation.

**Figure 2.**
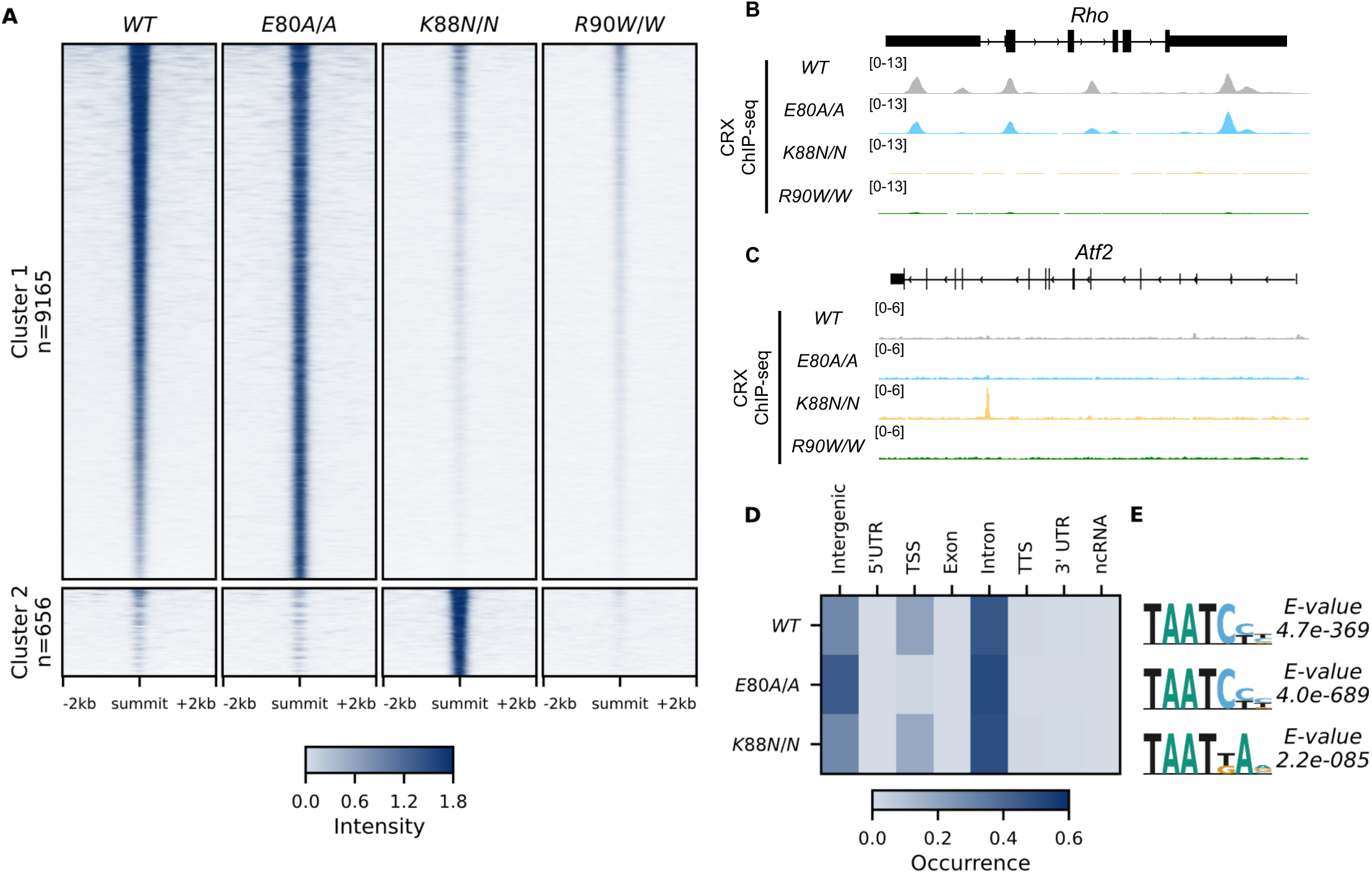
CRX E80A binds to WT sites while CRX K88N occupies novel genomic regions enriched for N88 HD motif *in vivo*. (A) Enrichment heat map depicting CRX ChIP-seq normalized reads centered at all possible CRX peaks ± 2kb, sorted by binding intensity in *WT* samples. Clusters were defined by hierarchical clustering of CRX binding intensity matrix from all genotypes (STAR Methods). (B-C) Genome browser representations of ChIP-seq normalized reads for different CRX species in P14 *WT* and mutant mouse retinas at *Rho* and *Atf2*. (D) Enrichment heatmap showing fraction of CRX ChIP-seq peaks fall in different genomic environments. (E) Logo representations of *de novo* found short HD motifs under CRX ChIP-seq peaks in *WT* and mutant mouse retinas with DREME *E-value* on the righ

**Figure 3.**
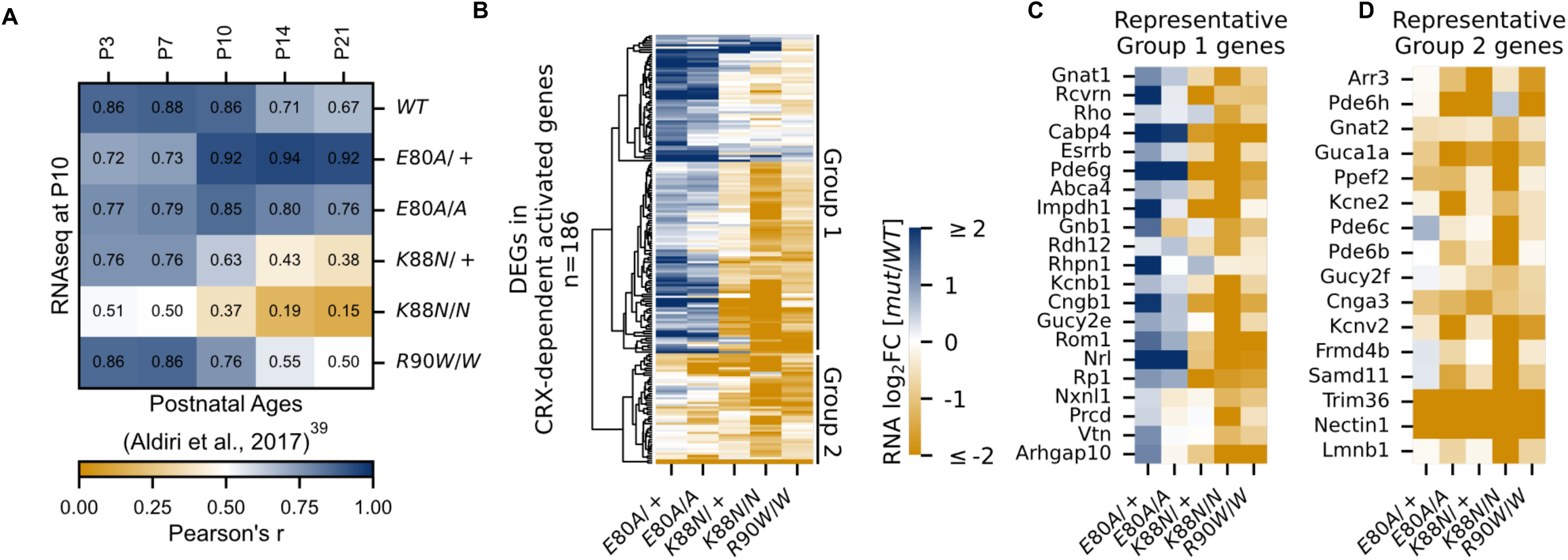
CRX-dependent activated genes affected in opposite directions in developing *Crx^E80A^* and *Crx^K88N^* mutant retinas. (A) Heat map showing sample-wise Pearson correlations of the expression of all CRX-dependent activated genes between P10 *WT* and HD mutant mouse retinas in this study (rows) with post-natal *WT* retinas from age P3 to P21 (columns, data from GSE87064). (B) Heat map showing the expression changes of DEGs in CRX-dependent activated gene set in HD mutant mouse retinas at P10. (C-D) Heat maps showing expression changes of selected photoreceptor genes from Group 1 and Group 2. Color scale identical to (B).

### E80A and K88N mutations affected the expression of CRX-dependent activated genes in opposite directions in a critical time window of photoreceptor differentiation

To understand how different CRX mutations affected gene expression at canonical and *de novo* binding sites and how these changes impair photoreceptor differentiation, we turned to bulk RNA-seq analysis from the developing retinas at P10. At P10, photoreceptors have started to differentiate, and the expression of many photoreceptor genes undergo exponential increase^39,40^. To focus on the most relevant expression changes, we first defined a set of genes that most likely depend on CRX activity nearby for expression (Figure S4A, Methods). Briefly, we associated each CRX ChIP-seq peak to the nearest gene, filtered only genes with nearby CRX ChIP-seq peaks, and further narrowed the list of genes to those significantly down-regulated in the loss of function mutant *Crx^R90W/W^*. Gene ontology (GO) analysis confirmed that this putative CRX-dependent gene set is associated with biological processes related to photoreceptor development and functions (Figure S4B). This set of putative CRX-dependent genes also showed developmental dependent gain in expression, consistent with CRX’s primary function as a transcriptional activator (Figure S4C). As a control, CRX-independent genes were constitutively expressed and largely involved in general cellular processes (Figures S4D-S4F). Therefore, the CRX-dependent gene set comprises genes important for photoreceptor differentiation and functional maturation and are dependent on CRX for activation. We denote these genes as “CRX-dependent activated genes” (*CRX-DAGs*).

Next, we sought to understand how each mutation affected photoreceptor differentiation. One way of measuring the progression of photoreceptor differentiation is to determine the similarity in *CRX-DAG* expression in a given sample with that of known developmental ages in *WT* control animals. We thus performed sample-wise correlation of *CRX-DAG* expression obtained in our P10 samples with a previously published RNA-seq dataset of normal mouse retinal development (Figure 3A)^39^. As expected, our P10 *WT* sample showed strong correlations with all developmental ages in the published *WT* control dataset. A stronger correlation with early ages (P3, P7, P10) and a weaker correlation with later ages (P14, P21) is also an indication of ongoing photoreceptor differentiation at P10. Unlike the *WT* sample, *Crx^E80A/+^* and *Crx^E80A/A^*samples both showed a stronger correlation with later developmental ages (P14, P21) but a weaker correlation with earlier postnatal ages (P3, P7). Since the *CRX-DAGs* are normally developmentally upregulated, this shift in correlation towards later developmental ages suggested that these genes were prematurely upregulated in the P10 *Crx^E80A^* mutant retinas. In contrast, *Crx^K88N/+^* and *Crx^K88N/N^* samples both showed a weaker correlation with all developmental ages when compared with *WT* samples in our dataset. This suggests that early photoreceptor differentiation was compromised in both *Crx^K88N^* mutants, consistent with their association with early-onset LCA^23^. Importantly, *Crx^R90W/W^*, also associated with LCA-like phenotype^19,22^, displayed strong correlation with earlier ages (P3, P7) similar to *WT*, but only showed moderate correlation with later ages (P14, P21). This suggests loss of CRX function at canonical binding sites does not affect the initiation of photoreceptor differentiation, but WT CRX activity at these sites is required to sustain differentiation. Since *CRX-DAG* expression was more severely affected in *Crx^K88N^* mutants than in *Crx^R90W/W^*, the photoreceptor differentiation deficits seen in the *Crx^K88N^* mutants cannot be explained solely by the loss of regulatory activity at canonical CRX binding sites. Overall, our sample-wise correlation analysis with normal retinal development dataset suggests that E80A and K88N mutations affected the expression CRX-dependent activated genes in opposite directions, implicating novel and distinct pathogenic mechanisms from the loss-of-function R90W mutation.

### *Crx^E80A^* retinas show up-regulation of rod genes but down-regulation of cone genes, underlying CoRD-like phenotype

Upon closer examination, we noted that not all *CRX-DAGs* were up-regulated in *Crx^E80A^* mutants (Figure 3B). Hierarchical clustering of all *CRX-DAGs* using expression changes revealed two major groups (Methods). In aggregate, when compared to *WT*, Group 1 genes were up-regulated in *Crx^E80A^* mutants while Group 2 genes were down-regulated. We noted genes indicative of the two photoreceptor subtypes, rods and cones, could partially define the two groups (Figures 3C and 3D). For example, *Esrrb*^41^ and *Nrl*^42^ in Group 1 are important regulators of rod differentiation. Other genes in Group 1 are components of the phototransduction cascade in rods, including *Rcvrn*^43^, *Rho*^44^, *Gnat1*^45,46^, *Pde6g*^47^, *Abca4*^48,49^, *Gnb1*^50^, *Rdh12*^51^, *Cngb1*^52^, *Rp1*^53,54^. Mis-regulations of many of these genes have been associated with diseases that affect rod development, function, and long-term survival. The increased activation of these genes likely underlies the stronger correlation with later developmental ages in *Crx^E80A+/^*and *Crx^E80A/A^* retinas (Figure 3A). In contrast, Group 2 genes, many down-regulated in *Crx^E80A^* mutants, were implicated in cone development and functions. For example, *Gnat2*^55,56^, *Pde6c*^57^, and *Pde6h*^58,59^ all act in the cone phototransduction cascade. Mis-regulation of these genes has also been implicated in different retinal dystrophies that primarily affect cone photoreceptors. Comparison of ChIP-seq signal revealed that peaks associated with Group 2 genes showed lower occupancy compared to Group 1 genes in the *Crx^E80A/A^* retinas (Mann-Whitney U test *p-value*: 9.51e-07) while no difference was observed in *WT* retinas (Mann-Whitney U test *p-value*: 0.541), suggesting loss of CRX activity likely underlies the down-regulation of Group 2 genes in the *Crx^E80A^* mutants (Figure S5A). Collectively, the selective down-regulation of cone genes in Group 2 may explain the CoRD-like phenotype in adult *Crx^E80A^*mutant mice described later.

Additionally, we noticed that a subset of genes not affected in *Crx^R90W/W^*(CRX-independent genes) were also down-regulated in *Crx^E80A^* mutants (Figures S5B-S5D). Among these genes were transcription regulators important for early photoreceptor development, such as *Ascl1*^60^, *Rax*^61^, *Sall3*^62^, and *Pias3*^63^. The down-regulation of these factors coincided with the up-regulation of mature rod genes in P10 *Crx^E80A^* retinas, suggesting that the E80A mutation might hamper the proper timing of photoreceptor differentiation.

### *Crx^K88N^* retinas display greater reduction of rod and cone genes than the loss-of-function mutants

*Crx^K88N^* retinas had the most severe gene expression changes among all mutants with down-regulation of both Group 1 (rod) and Group 2 (cone) genes (Figures 3B-3D). The heterozygous *Crx^K88N/+^* retina displayed a similar degree of expression reduction as homozygous *Crx^R90W/W^*, consistent with its association with dLCA^23^. Given the normal phenotype of heterozygous loss-of-function mutants - *Crx^+/-^*and *Crx^R90W/+^*^19^, these results suggest that mutant CRX K88N not only failed to activate WT target genes, but also functionally antagonized WT CRX regulatory activity in differentiating photoreceptors. This antagonism might be associated with ectopic CRX K88N activity when bound to regulatory regions with N88 HD DNA motifs (Figures 2A and 2E). In the absence of WT CRX, *Crx^K88N/N^* retina displayed a more severe expression reduction of *CRX-DAG* than *Crx^R90W/W^*, raising the possibility that CRX K88N also antagonized the activity of other transcriptional regulators important for photoreceptor differentiation. Supporting this possibility, a set of CRX-independent genes were also mis-regulated in both heterozygous and homozygous *Crx^K88N^* mutants. We noted that a number of these down-regulated genes are also involved in photoreceptor functional development (Figures S5B, S5E, and S5F). Overall, CRX K88N is associated with greater gene expression changes than other CRX HD mutants, which is likely attributed to ectopic regulatory activity.

### Both *E80A* and *K88N* mutants show compromised rod/cone terminal differentiation in young adults

Since *Crx^+/-^* and *Crx^R90W/+^* mutant mouse models showed a late-time recovery in photoreceptor gene expression and function^20^, we sought to determine the degree of photoreceptor differentiation in *Crx^E80A^* and *Crx^K88N^* mutants at P21. At this age, the normal retina has largely completed terminal differentiation with photoreceptor gene expression reaching a plateau^64^. However, when P21 CRX HD mutant retinas were examined for the expression of genes under the GO term *detection of light stimulus* (GO:0009583), which comprises genes in both rod and cone phototransduction cascades, many genes failed to reach *WT* levels, despite variable degrees of impact across different HD mutants (Figure 4A). *Crx^E80A^* mutants, in contrast to the increased rod gene expression at P10, displayed a deficiency in both cone and rod phototransduction genes at P21 (Figure 4B, Supplementary Table 6). This suggests that mutant CRX E80A transcriptional activity fails to sustain photoreceptor terminal differentiation and ultimately results in non-functional and severely affected photoreceptors. In comparison, *Crx^K88N^* mutants showed severely reduced expression of rod/cone phototransduction genes at both P10 and P21 (Figures 3B-3D and 4B).

**Figure 4.**
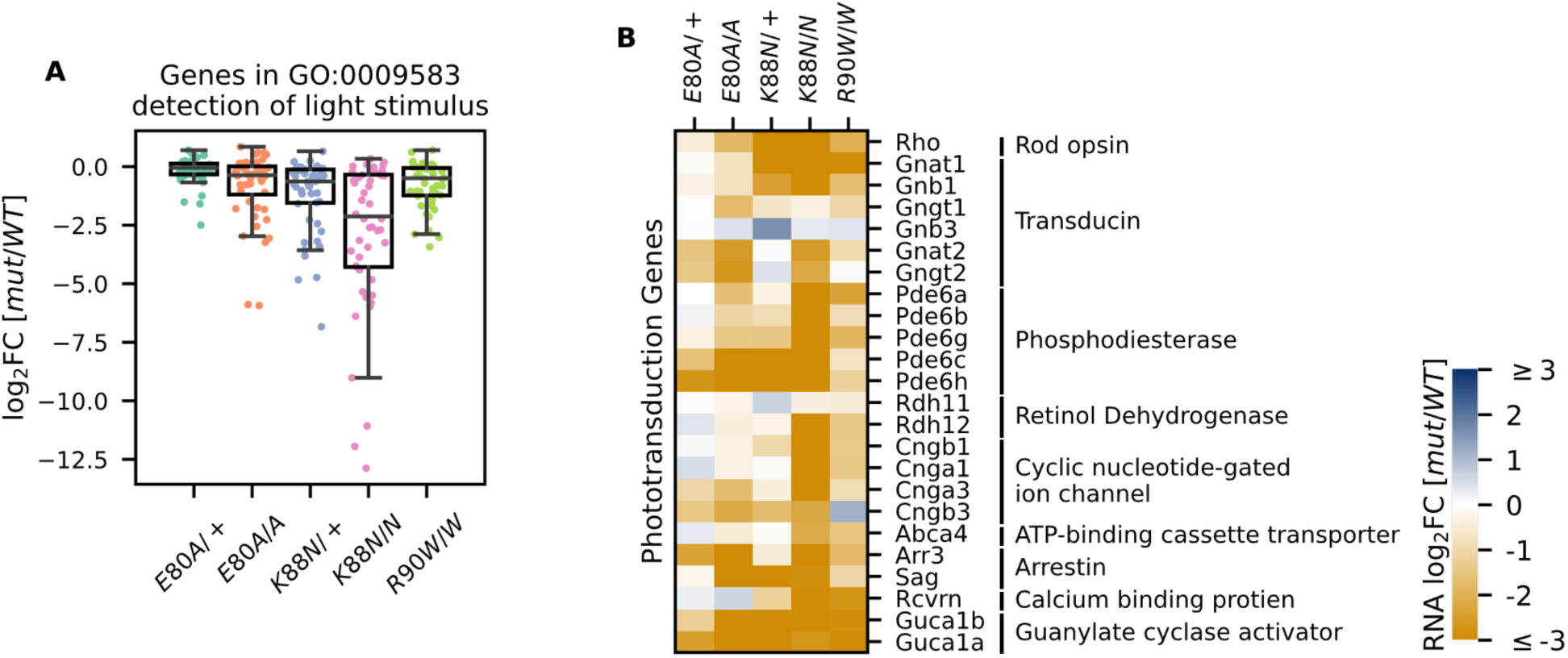
Photoreceptor genes important for phototransduction are down-regulated in all HD mutants. (A) Box plot showing that genes in the detection of light stimulus GO term were down-regulated and affected to various degrees in different adult (P21) HD mutant mouse retinas. (B) Heat map showing that expression of both cone and rod phototransduction genes were down-regulated in adult (P21) HD mutant mouse retinas. Annotation of rod and cone enrichment of each gene is in Supplementary Table S6. See Supplementary Figure S6 for the developmental expression dynamics of these genes.

To assess retinal morphology and photoreceptor subtype-specific gene expression at the cellular level, we performed immunohistochemistry analysis on P21 retinal sections. In *WT* animals, a hallmark of photoreceptor maturation is the outgrowth of photoreceptor outer segments (OS) filled with proteins necessary for the phototransduction. We thus performed hematoxylin and eosin (H&E) staining on P21 sagittal retinal sections to visualize changes in retinal layer organization, focusing on photoreceptor layers – outer nuclear layer (ONL) and OS. Compared to the well-organized ONL in *WT* retinas (Figure 5A), all mutants showed variable degrees of ONL disorganization, forming waves, whorls, and rosettes (Figures 5B-5E). The ONL disorganization was more severe in homozygotes than in heterozygotes for both mutations and *Crx^K88N^*mutants were more severely affected than *Crx^E80A^* mutants. Photoreceptor OS layer was formed in the *Crx^E80A/+^* retinas, but absent in *Crx^E80A/A^*, *Crx^K88N/+^*, and *Crx^K88N/N^* mutant retinas. Inner retinal layers, including inner plexiform layer (IPL) and ganglion cell layer (GCL) were not as severely affected as the outer retinal layers, supporting a model that the mutant morphological abnormalities largely originated from the diseased photoreceptors. These morphological abnormalities were distinct from the degenerative phenotypes of other *Crx* mutant models reported previously^19,65,66^.

**Figure 5.**
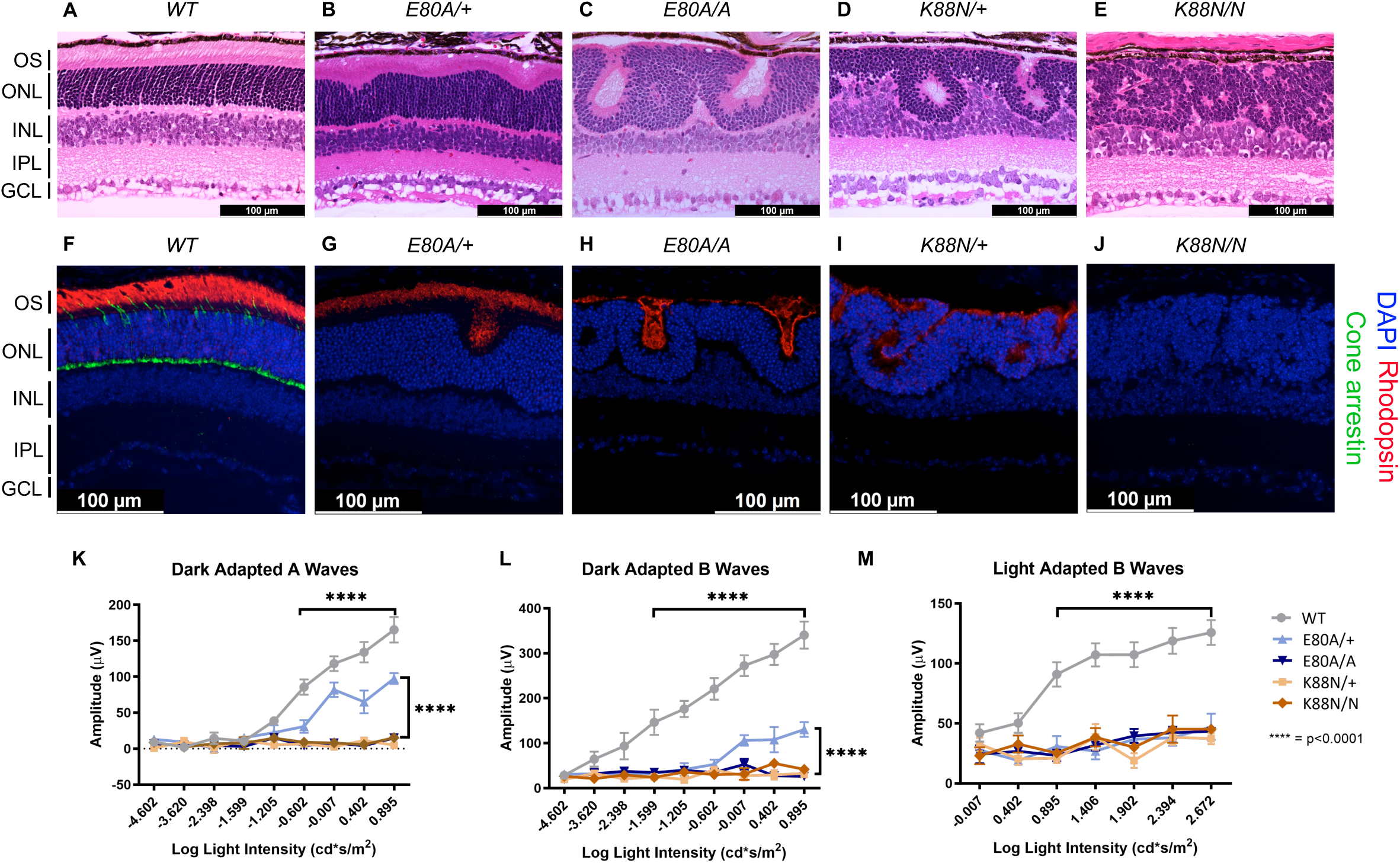
Only *Crx^E80A^*^/+^ retinas maintain photoreceptor OS and residual rod ERG response. (A-E) Hematoxylin-eosin (H&E) staining of P21 retina sections show that photoreceptor OS layer is absent in all mutant retinas except *Crx^E80A/+^*. OS: outer segment; ONL: outer nuclear layer; INL: inner nuclear layer; IPL: inner plexiform layer; GCL: ganglion cell layer. Scale bar, 100µm. (F-J) Rhodopsin (RHO, red) immunostaining is present in *Crx^E80A/+^*, *Crx^E80A/A^*, and *Crx^K88N/+^* retinas and absent in *Crx^K88N/N^* retina. Cone arrestin (mCAR, green) immunostaining is absent in all mutant retinas. Nuclei were visualized by DAPI staining (Blue). Scale bar, 100µm. (K-M) The electroretinogram responses (ERG) recorded from 1-month mice. Error bars represent the standard error of the mean (SEM, n ≥ 4). *p-value*: Two-way ANOVA and Tukey’s multiple comparisons. ****: p ≤ 0.0001. ns: >0.05.

Next, we sought to determine the expression of the rod-specific visual pigment rhodopsin (RHO) and cone arrestin (mCAR) in the P21 mouse retinas. In *WT* retinas, RHO is trafficked to the rod OS while mCAR is present in the cone OS and IS (inner segment), cell body, and synaptic terminals (Figure 5F). Unlike *WT* retina, all mutants lacked mCAR immunoreactivity (Figures 5G-5J), consistent with the loss of cone gene expression shown by RNAseq (Figure 4B). In *Crx^E80A/+^* retinas, RHO staining was localized to the OS layer; in *Crx^E80A/A^* and *Crx^K88N/+^*retinas, positive RHO staining was observed within the whorls and rosettes; in *Crx^K88N/N^* mutant retinas, RHO staining was completely absent. Importantly, we did not observe mis-localized RHO staining in the inner retinal layers (INL) suggesting that the developmental programs of other retinal cell types were not directly affected by E80A or K88N mutation. Overall, abnormalities in the cone/rod gene expression matched the corresponding human disease diagnosis^21,23^, and the phenotypic severity correlated with the degree of mis-regulation of CRX target genes in the corresponding RNAseq dataset. Thus, these results support a model that CRX HD mutation-mediated mis-regulation of gene expression disrupts photoreceptor terminal differentiation and leads to defects in retinal layer organization and OS formation.

### *E80A* and *K88N* mouse models show visual function deficits that recapitulate human diseases

To understand the consequences of disrupted photoreceptor differentiation on visual function, we measured electroretinogram (ERG) responses to light stimuli for *WT* and mutant mice at one month of age (Figures 5K-M). In response to incremental changes of light intensities, *WT* animals showed corresponding amplitude increases in dark adapted A-waves (rod signals) and B-waves (rod-evoked bipolar cell signals), as well as in light-adapted B-waves (cone-evoked bipolar cell signals). The three severe mutants, *Crx^E80A/A^*, *Crx^K88N/+^*, and *Crx^K88N/N^* had no detectable dark-adapted or light-adapted ERG responses, suggesting that these mice have no rod or cone function and are blind at young ages. The null ERG phenotype of the *Crx^K88N^* animals is consistent with the clinical LCA phenotype in humans^23^. In contrast, *Crx^E80A/+^* animals retained partial rod ERG responses as indicated by the reduced A-wave and B-wave amplitudes (Figures 5K and 5L). Yet, *Crx^E80A/+^* animals had no detectable cone ERG responses, which is consistent with the CoRD clinical phenotype in humans^21^ (Figure 5M). Taken together, the visual function impairment in each CRX HD mutant model, coincided with the morphological and molecular changes, suggesting that *Crx^E80A^* and *Crx^K88N^*mouse models recapitulate the corresponding human diseases.

### CRX E80A has increased transactivation activity and leads to precocious differentiation in *Crx^E80A^* retinas

Lastly, we asked what might be the molecular mechanism that causes the mis-regulation of photoreceptor genes in the mutant retinas. Previous studies have established that reporter assays with the *rhodopsin* promoter in HEK293T cells can measure changes of CRX transactivation activity and inform the mechanisms by which photoreceptor genes are mis-regulated in CRX mutant retinas^18^. We thus tested the transactivation activity of the three CRX HD mutants on the *pRho-Luc* reporter in HEK293T cells (Figure 6A, Methods). Consistent with published studies, R90W mutant had significantly reduced activity compared to WT CRX^18^. K88N mutant showed a similarly reduced activity as R90W, consistent with K88N’s loss of binding at canonical CRX sites and failure to activate *CRX-DAGs in vivo* (Figures 2A, 2B, 3A-3C). In contrast, E80A mutant, which binds to canonical CRX sties, showed significantly increased transactivation activity on the *Rho* promoter. This hyperactivity of E80A protein at *Rho* promoter correlates with the upregulation of *CRX-DAGs* in the mutant retinas at P10 (Figures 3A-3C).

**Figure 6.**
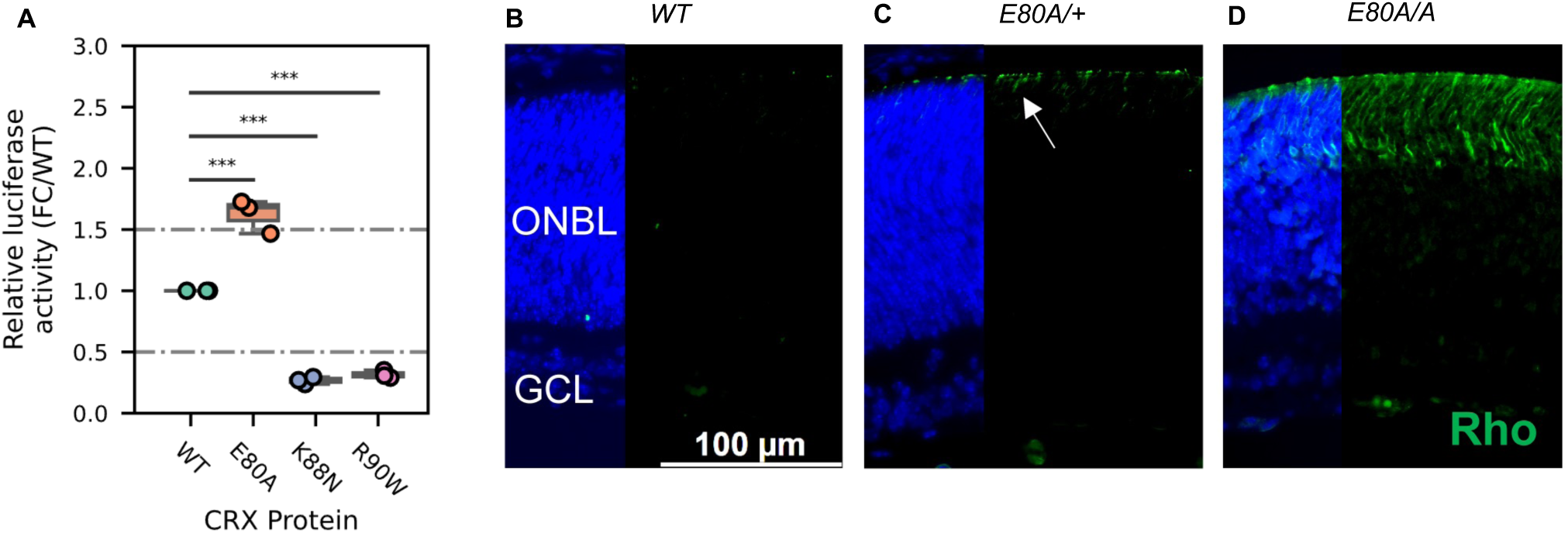
CRX E80A hyperactivity underlies precocious photoreceptor differentiation in *Crx^E80A^* retinas. (A) Boxplot showing luciferase reporter activities of different CRX variants. *P-values* for one-way ANOVA with Turkey honestly significant difference (HSD) test are indicated. *p-value*: ****: ≤0.0001, ***: ≤0.001, ns: >0.05. (B-D) Rhodopsin (RHO, green) immunostaining is absent in P3 *WT* retina but detected in *Crx^E80A/+^* and *Crx^E80A/A^* retinas. Nuclei are visualized by DAPI staining (Blue). Arrow indicates the sporadic RHO staining in *Crx^E80A/+^* sample. ONBL: outer neuroblast layer; GCL: ganglion cell layer. Scale bar, 100µm.

A transition to the next developmental stage usually requires the expression of important developmental genes passing an abundance threshold. Based on the hyperactivity model, photoreceptor genes are activated stronger in *Crx^E80A^* retinas and thus could reach the abundance threshold earlier. To determine the consequences of E80A hyperactivity on photoreceptor differentiation timing, we compared RHO protein expression during early postnatal retinal development using retinal section immunostaining (Figures 6B-6D). In *WT* retinas, most rods were born by P3 but had not differentiated (Figure 6B). Previous studies showed that RHO proteins were detected by IHC starting around P7 in *WT* retinas^64^. In comparison, both *Crx^E80A/+^* and *Crx^E80A/A^* retinas showed positive RHO staining at P3 (Figures 6C and 6D). RHO^+^ cells were largely seen in the outer portion of the ONBL layers in *Crx^E80A/+^* retinas, and strikingly spread throughout the large presumptive ONL layers in *Crx^E80A/A^* retinas. The detection of RHO protein in P3 *Crx^E80A^* mutant retinas indicates that photoreceptor differentiation program was precociously activated. Taken together, our results support a model that *E80A* and *K88N* mutations each perturbs CRX regulatory activity in a unique way, causes photoreceptor differentiation defects, and ultimately leads to distinct dominant disease phenotype that recapitulates human diseases.

## Discussion

Through molecular characterization of mutant proteins, transcriptome, and cellular profiling of developing mutant mouse retinas and ERG testing of adult retinas, we have identified two novel pathogenic mechanisms of CRX HD mutations, E80A and K88N, that are associated with dominant CoRD and dominant LCA in human^21,22^. Distinct from the previously characterized loss-of-function R90W mutation^19,20^, E80A and K88N mutations produce altered CRX proteins with gain of regulatory functions - CRX E80A is associated with increased transcriptional activity and CRX K88N has altered DNA binding specificity. Both CRX E80A and CRX K88N proteins impair photoreceptor gene expression, development and produce structural and functional deficits in knock-in mouse models, recapitulating human diseases^65^ (Figure 7). Thus, both target specificity and regulatory activity precision at the canonical CRX targets are essential for proper photoreceptor development and functional maturation.

**Figure 7.**
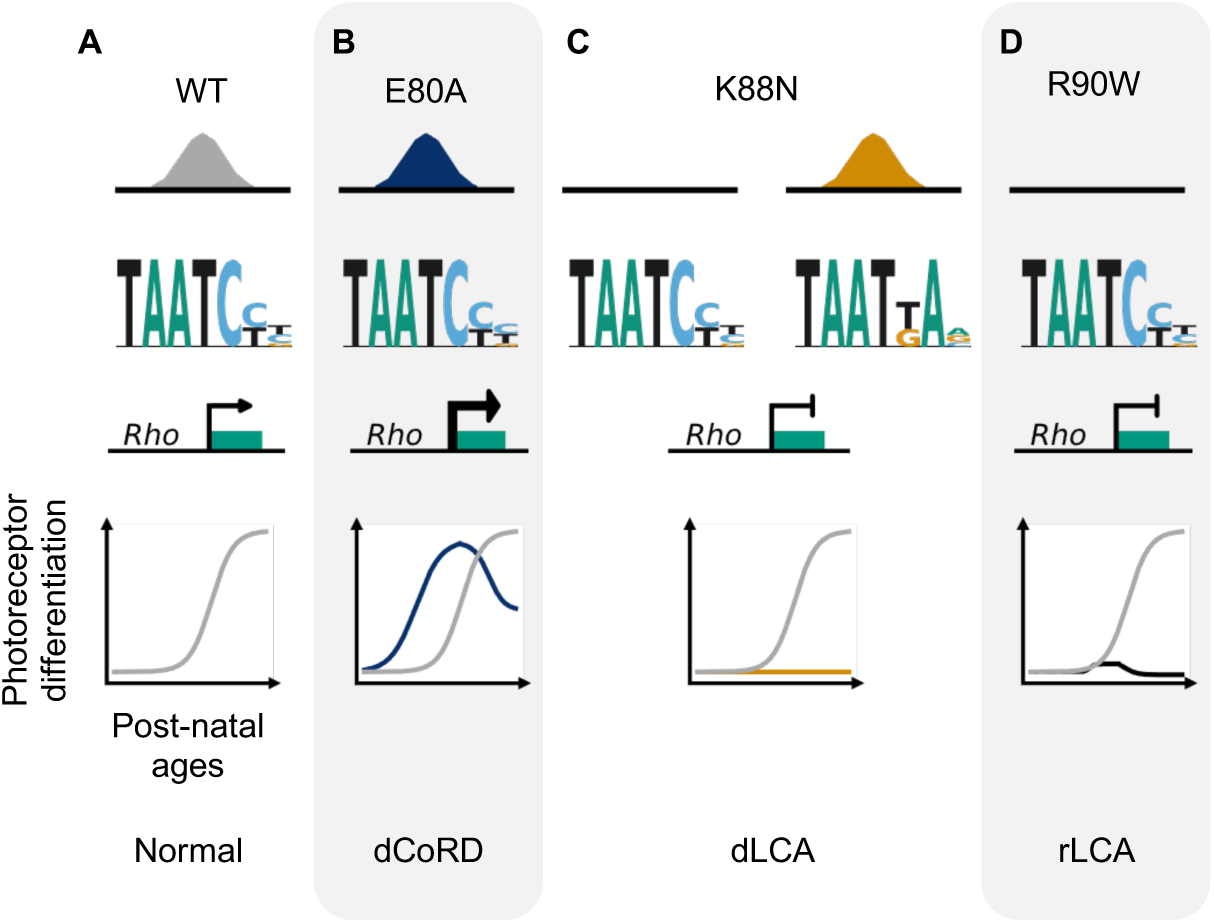
Missense mutations in CRX HD affect photoreceptor gene expression and leads to distinct retinal disease phenotypes through gain- and loss-of-function mechanisms.

Although associated with distinct disease phenotypes, the E80A, K88N, and R90W mutations are located very close to each other in the CRX HD recognition helix. Extensive biochemical and structural studies on the HD-DNA complexes afford important insights into how these mutations could affect CRX HD-DNA interactions differently. Most distinctively, CRX K88 residue, at HD50 position, is the major contributor to HD-DNA binding specificity, with lysine making favorable interactions with both guanines in the CRX consensus TAATCC binding site^67,68^. It is thus expected that K88N mutation drastically changes CRX DNA binding preference at the 3’ end of the HD motif, reminiscent of previous findings on novel HD DNA-binding specificity using bacterial one-hybrid (B1H) systems^31,69^. Supporting the importance of K88 residue mediated CRX target specificity in regulating photoreceptor development, *Crx^K88N^*retinas show more severe perturbations in photoreceptor gene expression and development than in loss-of-function mutant *Crx^R90W/W^*. *Crx^K88N/+^*and *Crx^K88N/N^* mice display the most severe photoreceptor morphological deficits observed in any *Crx* mouse models and show absence of visual functions in young adults. Thus, CRX target specificity is critical for photoreceptor development fidelity.

In developing *WT* mouse retinas, the HD motif preferred by HD TFs with a glutamine (Q) at HD50 position encodes quantitatively different activity than the CRX consensus suggesting functional difference between HD binding site variants^61,70^. It is likely that the severe *Crx^K88N^* phenotypes are attributed to both diminished activity at canonical CRX motifs and ectopic binding and transcriptional activity at N88 HD motifs. Since a functional copy of WT CRX is retained in *Crx^K88N/+^* retinas, the lack of WT activity alone cannot explain the severe developmental deficits. Alternatively, these results suggest involvement of additional regulatory mechanisms: CRX K88N activity at N88 HD motifs might (1) ectopically activate genes whose expression prevents the progression of development or inactivate genes required for development; (2) interfere with other HD TFs that also recognize N88 HD motifs; (3) lead to epigenetic alterations that antagonize normal CRX functions. Many other HD containing TFs are expressed in developing mouse retina, including OTX2, RAX, VSX2, PAX6, SIX3/6, and LHX family^6^. Different from CRX, these HD TFs are essential for gene regulation in retinal progenitor cells and/or in other retinal cell lineages. Alteration of CRX DNA binding specificity could mis-regulate genes originally targeted by these HD TFs and lead to severe perturbations in the retinal gene regulatory networks (GRNs). To date, most studies have focused on CRX activity at cis-regulatory sequences enriched for the WT CRX consensus motifs^71,72^. Systematic comparison of regulatory activity at N88 HD motifs and WT consensus in the context of photoreceptor development in both *WT* and mutant retinas would be needed to substantiate the impact of mutant CRX K88N activity at different HD motifs. These experiments will also help clarify the pathogenic mechanisms in the *Crx^K88N^*models and extend our knowledge of CRX HD mediated regulatory grammar during photoreceptor development.

Different from CRX K88, the E80 and R90 residues, although the most common residues at HD42 and HD52 positions respectively, do not contact DNA directly and thus lacked in-depth investigations in prior studies. CRX R90 residue has been suggested to confer additional stability for the homeodomain fold besides the core residues and make contacts with the DNA backbone through bases in the TAAT core motif^67,68^. Substitution of the basic R90 residue with a bulky, neutral tryptophan (W) potentially reduces overall CRX HD stability which in term reduces CRX HD DNA-binding affinity without affecting its binding preference. The potential reduction in CRX HD-DNA complex stability is in line with our observation that R90W mutation abolishes CRX binding across the genome resulting in global loss of CRX target gene activation. It also explains the association of CRX R90W mutation with recessive loss-of-function LCA phenotypes in human and mouse^22,65^.

Structural studies suggest that CRX E80 residue plays a role in stabilizing the HD-DNA binding complex through intramolecular interactions with other HD residues^29,68^, yet functional validations await further experiments. E80A mutation, replacing glutamic acid (E), which is acidic and polar, with alanine (A), which is neutral and non-polar, could render HD-DNA interactions more promiscuous as reflected in overall reduced magnitude of CRX E80A HD specificity (Figure 1I). Regulatory sequences of many photoreceptor genes contain both consensus and non-consensus CRX motifs^14,15,18,35^. It is likely that the more promiscuous CRX E80A-DNA interaction increases the likelihood of non-consensus CRX HD motifs being bound and activated resulting in overall increased transcriptional output (hyperactivity) as seen both in luciferase assays and in developing *Crx^E80A^* mouse retinas. The promiscuous TF-DNA-binding associated hyperactivity phenomenon has also been observed in a dominant disease mouse model harboring a missense mutation in the zinc finger TF Krüppel-like factor-1 (KLF1)^73^. Yet, in adult *Crx^E80A^* retinas, photoreceptor terminal differentiation is impaired, resulting in disrupted retinal morphology and defective visual functions. Photoreceptor differentiation is programmed via sequential and concerted gene expression programs within a defined time window^74,75^. One explanation for adult *Crx^E80A^* phenotypes is that CRX E80A hyperactivity precociously activates later stage genes in the absence of proper nuclear context and/or subcellular structures, which in turn negatively impacts early events in photoreceptor differentiation. These observations underscore the importance of precisely tuned CRX-mediated transcriptional activity during photoreceptor development.

Although a global increase in expression is expected in *Crx^E80A^* retinas based on the hyperactivity model, a subset of CRX dependent activated genes implicated in cone photoreceptor development and functions is down-regulated in both differentiating and mature mutant retinas. While cones undergo terminal differentiation to develop into cone subtypes – M- or S-cones – in a similar postnatal window as rods, they were born prenatally in mice within an earlier time window than rods. At an early postnatal age, cells expressing RXRγ, a ligand-dependent nuclear hormone receptor normally expressed in developing cones^60,76–78^, were observed in *Crx^E80A^* retinas, suggesting cone photoreceptors were born in these mutant retinas (Figure S7A). One model for lack of cone markers in adult *Crx^E80A^* retinas is that CRX E80A improperly activates later stage cone genes at a much earlier time window, disrupting cone terminal differentiation. Supporting this model, CRX E80A also hyperactivates the *S-cone opsin* promoter (Figure S7B). An alternative model is that cones might be more sensitive to perturbations in CRX activity. It is known that cones depend on a different repertoire of TFs than rods for subtype terminal differentiation^60^. It is possible that cone TFs respond differently to mutant CRX E80A hyperactivity, leading to the distinct expression changes in *Crx^E80A^* mutant retains. It is important to note that different point mutations at CRX E80 residue have been reported in dominant CoRD cases (ClinVar VCV000865803.1, VCV000007416.7, VCV000099599.6), emphasizing the importance of residue CRX E80 in regulating cone photoreceptor development. Since cones only make up a very small portion (3%) of photoreceptors in mouse retinas^79^, quantitative characterization of CRX E80A molecular functions in a cone dominant retina warrants further study to understand its selective effect on the cone differentiation program and help elucidate WT CRX regulatory principles in early photoreceptor development.

Given that the spatial structures and HD-DNA contact models of HD proteins are evolutionarily conserved, our study of CRX provides valuable molecular insights for HD mutations implicated in other diseases. For example, *p.E79K* substitution (corresponds to CRX E80) in OTX2 HD is associated with dominant early-onset retinal dystrophy^80^, heterozygous *p.R89G* (corresponds to CRX R90) mutation in OTX2 HD causes severe ocular malformations^81^, and missense mutations of the CRX K88 and R90 homologous residues in PITX2 HD are associated with dominant Rieger syndrome^82^. It is likely that these mutations affect HD activity in similar ways as observed in CRX, and the exact disease manifestation is determined by cell-type or tissue specific mechanisms. The retina is readily accessible, and a broad range of molecular tools are available for *ex vivo* and *in vivo* manipulations. We believe that CRX is an ideal model to study the pathogenic mechanisms of HD mutations and to test therapeutic regimens, which would ultimately benefit the study of HD TFs and their associated diseases in other tissues and organs.

One limitation of this work is that effects of E80A and K88N mutations on CRX HD-DNA interactions have been evaluated at monomeric HD motifs and with homogenous protein species both *in vitro* and *in vivo*. Further evaluation of WT and mutant CRX binding at dimeric motifs will be desirable, since selected dimeric HD motifs are known to mediate HD TF interactions to ensure gene expression fidelity during development^83,84^. Relatedly, we also need to address how CRX WT and mutant E80A or K88N proteins interact at HD binding motifs – whether they cooperate or compete with each other, whether these interactions are HD motif sequence-dependent, and how gene expression is impacted by CRX cooperativity or competition. While CRX HD mediates both TF-DNA interactions and protein-protein interactions, evaluation of how E80A and K88N mutations impact CRX interaction with other important photoreceptor TFs and how perturbations in these interactions lead to disease phenotypes warrant further study.

Collectively, our findings support a unifying model in which precise CRX interaction with cis-regulatory sequences is essential for gene expression and functional maturation during photoreceptor development. Disease-associated mutations in CRX have been classified into two main groups – insertion/deletion-derived frameshift mutations in the activation domain (AD) and missense mutations in the homeodomain^65^. Prior biochemical and mouse model studies of the first group have established that AD-truncated mutant proteins abolish CRX transcriptional activity and functionally interfere with the *WT* allele. As a result, the mutant retinas fail to activate or maintain robust cone/rod gene expression, resulting in incomplete photoreceptor differentiation and ultimately rapid degeneration of immature photoreceptors^16,19^. In this study, we demonstrate that missense mutations in the CRX HD, by either a loss- or gain-of-function mechanism, alter CRX target specificity and/or CRX transactivation activity. These biochemical property changes impair CRX-mediated transcriptional regulation *in vivo* and lead to distinct morphological and functional deficits (Figures 7A-7D). Despite the difference in molecular mechanisms, both *Crx^E80A^* and *Crx^K88N^* mouse models develop whorls and rosettes in the ONL by P21, which are not observed in degenerative CRX mouse models^19^, suggesting distinct pathogenic mechanisms. Future cellular biology studies are needed to understand the formation mechanisms of these unique cellular phenotypes (ONL disorganization) and their impacts on the function and survival of photoreceptors and other retinal cell types over development.

Our study here also emphasizes the importance of tailoring gene therapy regimens to tackle individual pathogenic mechanisms. For instance, while supplementing WT CRX might be sufficient to rescue a hypomorphic/loss-of-function mutant, simultaneous elimination of a gain-of-function *CRX* product would be necessary to rescue dominant mutants, as exemplified in a recent report of allele-specific gene editing to rescue dominant CRX-associated LCA7 phenotypes in a retinal organoid model^85^. We believe that this principle also applies to other dominant neurological diseases. Additionally, with the refinement of the CRX mechanistic model, when new disease mutations are identified, genetic counsellors can now provide more informed predictions of disease progression and future visual deficits. This information is important for individuals to be psychologically prepared and seek necessary assistance to improve their quality of life.

## Author contributions

S. Chen conceived and supervised the study. S. Chen, P. Ruzycki, C. Sun and Y. Zheng designed the experiments. Y. Zheng performed Spec-seq and luciferase experiments; X. Zhang performed CRX ChIP-seq experiments; C. Sun performed immunochemistry and ERG experiments. Y. Zheng analyzed Spec-seq, CRX ChIP-seq, RNA-seq, and luciferase data; P. Ruzycki assisted in CRX ChIP-seq and RNA-seq data analysis; C. Sun analyzed immunochemistry and ERG data. Y. Zheng wrote the original draft; S. Chen, P. Ruzycki, C. Sun, and Y. Zheng revised the manuscript. All authors read and approved the final manuscript.

## Acknowledgements

We thank Mingyan Yang and Guangyi Ling for technical assistance, Susan Penrose and Mike Casey from the Molecular Genetics Service Core for generating E80A and K88N mutation knock-in mice lines, Inez Oh for RNA-seq sample collection and processing, and J. Hoisington-Lopez and M. Crosby from DNA Sequencing Innovation Lab at the Center for Genome Sciences & Systems Biology for sequencing assistance. This work was supported by NIH grants EY012543 (to S. Chen), EY032136 (to S. Chen), EY002687 (to WU-DOVS), and the Stein Innovation Award (to SC) and unrestricted funds (to WU-DOVS) from Research to Prevent Blindness. We also thank Mr. Artur Widlak for the generous gift from Widłak Family CRX Research Fund.

## Inclusion and diversity

We support inclusive, diverse, and equitable conduct of research.

## Methods

### Resource availability

#### Lead contact

Further information and requests for resources and reagents should be directed to and will be fulfilled by the lead contact Shiming Chen.

#### Materials Availability

▪ All unique/stable reagents generated in this study are available from the lead contact with a completed materials transfer agreement.

#### Data and Code Availability

▪ The raw sequencing data and processed data generated in this study have been deposited at NCBI GEO.
▪ This paper does not report original code.
▪ Any additional information required to reanalyze the data reported in this paper is available from the lead contact upon request.

### Animal study and sample collection

#### Ethics statement

All procedures involving mice were approved by the Animal Studies Committee of Washington University in St. Louis and performed under Protocol 21-0414 (to SC). Experiments were carried out in strict accordance with recommendations in the Guide for the Care and Use of Laboratory Animals of the National Institutes of Health (Bethesda, MD), the Washington University Policy on the Use of Animals in Research; and the Guidelines for the Use of Animals in Visual Research of the Association for Research in Ophthalmology and Visual Sciences. Every effort was made to minimize the animals’ suffering, anxiety and discomfort.

#### Mutation knock-in mouse model generation

CRISPR/Cas9 based genome editing was performed to generate the *Crx^E80A^* and *Crx^K88N^* mice as previously described^86^. The Cas9 guide RNAs (gRNA) were designed based on proximity to the target amino acid and was synthesized using the MEGAshortscript™ T7 Transcription Kit (Thermo Fisher Scientific, Waltham, MA). The gRNAs were subsequently tested for cutting efficiency in cell culture by the Washington University Genome Engineering and iPSC Center. The validated gRNA and Cas9 protein were then microinjected into the pronuclei of C57Bl/6J-0.5-dpc (days post coitum) zygotes along with the donor DNA, a 190-bp single stranded oligodeoxynucleotide (ssODN) carrying either the *c.239A>G* substitution for *p.E80A* mutation or the *c.264G>T* substitution for *p.K88N* mutation^87^. Embryos were then transferred into the oviduct of pseudo-pregnant female. Pups were generally delivered ∼20 days after microinjection. Tissues from 10-day postnatal (P10) pups were collected by toe biopsy/tail for identification of the targeted allele by restriction digest (HinfI) of PCR amplified DNA first and then confirmed by Sanger sequencing (Genewiz).

Founders carrying the correct alleles were then bred with wild-type C57BL/6J mice (Jackson Laboratories, Bar Harbor, ME, Strain #000664) to confirm transmission. All experimental animals used were backcrossed at least 10 generations. Genotyping of mutation knock-in mice follows cycling conditions: 95°C for 2 min, 94°C for 30 sec, 60°C for 30 sec, 68°C for 60 sec, repeat steps 2-4 for 34 cycles, 68°C for 7 min, and hold at 4°C. After PCR reaction, the amplified DNA fragments wee digested with HinfI. Sequences of gRNAs, ssODNs and genotyping primers can be found in Supplementary Table S3. A representative DNA gel of HinfI digested genotyping DNA fragments can be found at Figure S3B.

#### RNA-seq sample collection and library preparation

For each genotype, three biological replicates, two retinas per replicate from one male and one female mouse were analyzed. All retinas were processed for RNA simultaneously using TRIzol™ Reagent (Invitrogen, Waltham, MA) following the manufacturer’s protocol. The quantity and quality of the RNA was assayed using Bioanalyzer (Agilent, Santa Clara, CA). Samples with a minimum RNA integrity number (RIN) score of 8.0 were then selected for library construction as previously described^20^.

#### Chromatin immunoprecipitation and library preparation

CRX chromatin immunoprecipitation was performed as previously published^88^. Briefly, pooled nuclear extracts from 6 retinae were cross-linked with formaldehyde prior to immunoprecipitation with anti-CRX antibody A-9 (#sc-377138, Santa Cruz Biotechnology, Dallas, TX). Input controls were included as background. The libraries were prepared following the standard ChIP-seq protocol^89^. The quantity and quality of the ChIP-seq libraries was assayed using Bioanalyzer (Agilent, Santa Clara, CA) prior to sequencing.

#### ERG and statistical analyses

ERGs were performed on 1-month-old mice using UTAS-E3000 Visual Electrodiagnostic System (LKC Technologies Inc., MD). Mice were dark-adapted overnight prior to the tests. Mouse body temperature was kept at 37 ± 0.5 °C during the tests. Pupils were dilated with 1% atropine sulfate solution (Bausch and Lomb). Platinum 2.0 mm loop electrodes were placed on the cornea of each eye. A reference electrode was inserted under the skin of the mouse’s head and a ground electrode was placed under the skin near mouse’s tail. Retinal response to full-field light flashes (10 μs) of increasing intensity were recorded; maximum flash intensity for dark-adapted testing was 0.895 cd*s/m2. Following dark adapted tests, mice were light-adapted under light condition (about 29.2 cd/mm) for 10 minutes and exposed to 10 μs light flashes of increasing intensity; maximum flash intensity for light-adapted testing was 2.672 cd*s/m2. ERG responses of biological replicates were recorded, averaged and analyzed using Graphpad Prism 8 (GraphPad Software, CA). The mean peak amplitudes of dark-adapted A and B waves and light-adapted B waves were plotted against log values of light intensities (cd*s/m2). The statistics were obtained by two-way ANOVA with multiple pairwise comparisons (Tukey’s).

#### Histology and immunohistology chemistry

Enucleated eyes were fixed at 4°C overnight for paraffin embedded sections. Each retinal cross-section was cut 5 microns thick on a microtome. Hematoxylin and Eosin (H&E) staining was performed to examine retinal morphology. For IHC staining, sections firstly went through antigen retrieval with citrate buffer, and blocked with a blocking buffer of 5% donkey serum, 1% BSA, 0.1% Triton X-100 in 1x PBS (pH-7.4) for 1 h. Sections were then incubated with primary antibodies at 4°C overnight. Sections were washed with 1x PBS containing 0.01% Triton X-100 (PBST) for 30 min, and then incubated with specific secondary antibodies for 1 h. Primary and secondary antibodies were applied with optimal dilution ratios. All slides were mounted with VECTASHIELD®HardSet™ Antifade Mounting Medium with DAPI (Vector Laboratories, Inc., CA). All images were taken on a Leica DB5500 microscope. All images were acquired at 1000 µm from ONH for ≥ P21 samples and at 500 µm from ONH for P0, P3, P10 samples.

### Biochemistry

#### Protein expression

Expression plasmids for GST-WT, E80A, and R90W HDs were published previously^18^. Plasmid for GST-K88N HD was generated by site-directed mutagenesis from the pGEX4T2-CRX WT HD backbone. *In vivo* protein expression and purification was done as previously described^18^. Briefly, 0.05mM IPTG was added to *E. coli* BL-21 (DE3) cell cultures containing different CRX HD constructs at OD_600_ = 0.6. The cultures were incubated for 2 hours or until OD_600_ = 2.0 at 34°C and the cells were collected by centrifugation at 6000 rpm and 4°C for 15 minutes. Cell pellets were resuspended in 1x PBS (Corning, Corning, NY) and then lysed by sonication. 5mM DTT (Bio-Rad Laboratories, Inc., Hercules, CA) and 1% Triton X-100 (MilliporeSigma, Burlington, MA) was then added, and the mixtures were incubated with gentle shaking at 4°C for 30 minutes to maximize protein extraction. The separation of proteins from the cellular debris were then performed by centrifugation at 15 000 rpm for 10 minutes and filtered through a 0.45μm membrane. Glutathione Sepharose 4B resin (Cytiva, Marlborough, MA) was first equilibrated with PBS before adding to the supernatant. 5x Halt™ Protease Inhibitor Cocktail and PMSF was added to minimize degradation. The mixtures were incubated with gentle shaking at 4°C overnight before loading on GST Spintrap™ columns (Cytiva, Marlborough, MA). The peptides were eluted following the manufacturer’s protocol and buffer exchanged into CRX binding buffer^90^ using Amicon centrifugal filters (MilliporeSigma, Burlington, MA). The protein stock was supplemented with 10% glycerol before aliquoted and stored at -80°C.

#### Protein quantification and visualization

The size and integrity of purified GST-CRX HDs were visualized with a native 12% Tris-Glycine SDS-PAGE gel in the absence any reducing agent. Protein concentration was measured by NanoDrop Oneᶜ Microvolume UV-Vis Spectrophotometers (ThermoFisher Scientific, Waltham, MA) and calculated using the equation: C = (1.55 * A_280_) – (0.76 * A_260_), where C is the concentration of the protein in mg/ml, A_280_ and A_260_ are the absorbance of protein samples at 280nm and 260nm respectively^91^. The protein concentrations obtained with this method were comparable with BCA protein quantification assays.

#### Spec-seq library synthesis and purification

Single-stranded Spec-seq library templates and IRDye 700-labeled reverse complement primers (Supplementary Table S1) were ordered directly from Integrated DNA Technologies (IDT, Coralville, Iowa). The synthesis and purification of the double-stranded libraries followed previously published protocols^33,34,91^. Briefly, 100 pmol of template oligos and 125 pmol IRDye 700-labeled reverse complement primer F1 were mixed in Phusion® High-Fidelity PCR Master Mix (NEB, Ipswich, MA). A 15s denaturing at 95°C following a 10-minute extension at 52°C afforded duplex DNAs. Subsequently, the mixture was treated with 1ul Exonuclease I (NEB, Ipswich, MA) to remove excess ssDNA. The libraries were purified by MinElute PCR Purification Kit (QIAGEN, Hilden, Germany) and eluted in molecular biology graded water (Corning, Corning, NY).

#### EMSA and sample preparation for sequencing

The protein-DNA binding reactions was done in 1x CRX binding buffer (60mM KCl, 25mM HEPES, 5% glycerol, 1mM DTT)^90^. A fixed amount (Supplementary Table S1) of IRDye-labelled DNA libraries were incubated on ice for 30 minutes with varying concentrations of wild type or mutant peptides in 20μl reaction volume. The reaction mixtures were run at 4°C in native 12% Tris-Glycine PAGE gel (Invitrogen, Waltham, MA) at 160V for 40min. The IRDye-labeled DNA fragments in the bound and unbound fractions were visualized by Odyssey® CLx and Fc Imaging Systems (LI-COR, Inc., Lincoln, NE). The visible bands were excised from the gels and DNAs were extracted with acrylamide extraction buffer (100mM NH_4_OAc, 10mM Mg(OAc)_2_, 0.1% SDS) then purified with MinElute PCR Purification Kit (QIAGEN, Hilden, Germany). The DNAs were amplified, barcoded by indexed Illumina primers. All indexed libraries were then pooled and sequenced on a single 1×50bp Miseq run at DNA Sequencing Innovation Lab at the Center for Genome Sciences & Systems Biology (CGS&SB, WashU).

#### qRT-PCR

For each replicate, RNA from 2 retinae of a mouse was extracted using the NucleoSpin® RNA kits (Takara Bio USA, Inc., San Jose, CA). RNA sample concentration and quality was determined with NanoDrop Oneᶜ Microvolume UV-Vis Spectrophotometers (ThermoFisher Scientific, Waltham, MA). 1 μg of RNA was used for cDNA synthesis with iScript cDNA Synthesis Kits (BioRad, Hercules, CA) in a 20μl reaction volume. Primers used in this study were listed in Supplementary Table S2. qRT-PCR reactions were assembled using SsoFast™ EvaGreen® Supermix with Low ROX (Bio-Rad Laboratories, Inc., Hercules, CA) following manufacturer’s protocol. Data was obtained from Bio-Rad CFX96 Thermal Cycler following a three-step protocol: 1 cycle of 95°C 3 min, 40 cycles of 95°C 10 sec and 60°C 30 sec. Data was exported and further processed with customized python script.

#### Western blot

Experiments were performed using two biological replicates with two retinas for each replicate. Nuclear extracts were prepared using the NE-PER Nuclear and Cytoplasmic Extraction Reagents (Thermo Fisher Scientific, Waltham, MA) following manufacturer’s instructions. 1x Roche cOmplete™ Mini Protease Inhibitor Cocktail (MilliporeSigma, Burlington, MA) was supplemented in all extraction reagents. 5mM of DTT was added immediately before sample denaturing and protein was separated by running on Invitrogen NuPAGE™ Novex 4-12% Bis-Tris MiniGels (Invitrogen, Waltham, MA). Membrane transfer was done with the Blot™ mini blot module (Invitrogen, Waltham, MA) following the manufacturer’s protocol. Membrane was probed with mouse monoclonal anti-CRX antibody M02 (1:1 000, Abnova Corp., Taipei City, Taiwan) and rabbit polyclonal anti-HDAC1 antibody H51 (1:1 000, Santa Cruz Biotechnology, Dallas, TX), visualized with IRDye® 680RD goat anti-rabbit IgG and IRDye® 800CW goat anti-mouse IgG secondary antibodies (1:10 000, LI-COR, Inc., Lincoln, NE). The membrane was then imaged using Odyssey® CLx and Fc Imaging Systems (LI-COR, Inc., Lincoln, NE).

#### Transient transfection luciferase reporter assays

HEK293T cells were cultured in Dulbecco’s Modified Eagle Medium supplemented with 10% fetal bovine serum (FBS) and Penicillin-Streptomycin following the manufacturer’s protocol. Cells were transfected with calcium phosphate transfection protocol in 6-well plates as previously described^18,19^. Experimental plasmids and usage amount are described in Supplementary Table S1. Typically, 48 hours after transfection, cells were harvested, digested and assayed for luciferase activity using Dual-Luciferase Reporter Assay System (Promega, Madison, WI) following the manufacturer’s protocol. Data were collected using TD-20/20 Luminometer (Turner Designs, East Lyme, CT) and further processed with customized python scripts.

### Data analysis

#### Homeodomain sequence alignment

The full-length protein sequences for the selected TFs were first aligned with Clustal Omega (EMBL-EBI, UK). Aligned sequences of the third homeodomain helix were then extracted to generate Figure 1B using Jalview (v2.11.1.7). A list of the accession numbers for the selected TFs can be found in Supplementary Table S1.

#### Determination of relative binding affinity with Spec-seq

For a biomolecular interaction between a protein *P* and a particular DNA sequence, *S_i_*, the interaction can be diagrammed as:

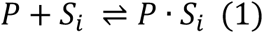

where *p* · *i_i_* refers to the protein-DNA complex. The affinity of the protein *P* to sequence *S_i_* is defined as the association constant *K_A_*, or its reciprocal, the dissociation constant *K_D_*. The *K_A_* of the protein-DNA interaction is determined by measuring the equilibrium concentrations of each reactant and the complex:

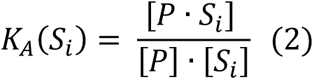

where […] refers to concentrations. As in a typical Spec-seq experiment, thousands of different DNA sequences compete for the same pool of proteins, their relative binding affinities (the ratio of their *K_A_*) can be determined by measuring the concentrations of each sequence in the bound and unbound fractions without measuring the free protein concentrations, which is often the most difficult to measure accurately:

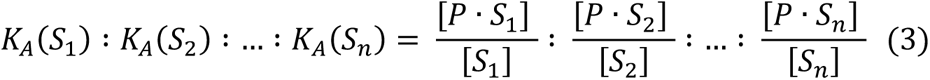

In a binding reaction involving TF and a library of DNAs, the concentration of bound and unbound species are directly proportional to the number of individual DNA molecules in each fraction which can be obtained directly from sequencing data. With enough counts in each fraction, we can accurately estimate the ratios of concentrations from counts with the relationship:

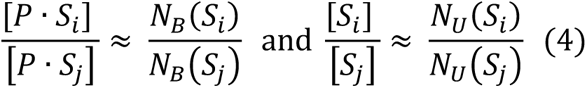

where *N_U_* denotes counts in the unbound fraction and *N_B_* denotes counts in the bound fraction. Therefore, the binding affinity of a sequence variant *S_x_* relative to the reference sequence *S_ref_* can be calculated by:

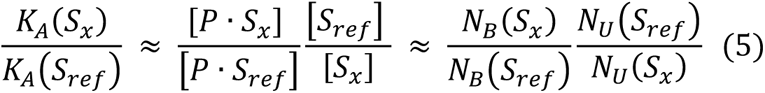

The natural logarithms of these ratios are the relative binding free energies in the units of *kcal/mol*. The relative free energy of the reference site for each CRX HD was set to 0.

#### Spec-seq data analysis and energy logo visualization

The sequencing results were first filtered and sorted based on conserved regions and barcodes. Reads with any mismatch in the conserved regions were discarded prior to further analysis as described previously^32,34,91^. The ratio of individual sequence in bound and unbound reads was calculated as a measurement of relative binding affinity (Equation. 5) compared to the consensus sequence. The relative binding energy was then derived from the natural logarithm of the relative binding affinity and represented in *kcal/mol* units.

For wild type CRX HD and all the mutants, the energy weight matrices (ePWMs) were generated based on the regression of the TF’s binding energy to its reference sequence. Only sequences within two mismatches to the reference were used to generate the ePWMs. Energy logos were generated from ePWMs after normalizing the sum of energy on each position to 0 and the negative energy values were plotted such that preferred bases are on top. The sequence logos were generated from ePWMs with python package logomaker (v0.8). The ePWMs for all CRX HDs are listed in Supplementary Table S3.

#### ChIP-seq data analysis

2×150bp reads from Illumina NovaSeq were obtained for all samples with a minimum depth of 22M reads at Novogene (Beijing, China). For each sample, reads from two sequencing lanes were first concatenated and run through Trim Galore (v0.6.1)^92^ to remove adapter sequences and then QC by FastQC (v0.11.5)^93^. The trimmed reads were then mapped to the mm10 genome using Bowtie2 (v 2.3.4.1)^94^ with parameters -X 2000 --very-sensitive. Only uniquely mapped and properly paired reads were retained with samtools (v1.9)^95^ with parameters -f 0×2 -q 30. Mitochondria reads were removed with samtools (v1.9)^95^. Duplicated reads were marked and removed with Picard (v2.21.4)^96^. Last, reads mapped to the mm10 blacklist regions were removed by bedtools (v2.27.1)^97^ by intersect -v. bigWig files were generated with deeptools (v3.0.0)^98^ with command bamCoverage --binSize 10 -e -- normalizeUsing CPM and visualized on IGV Web App^99^. For each genotype, an average binding intensity bigWig file from two replicates was generated with deeptools (v3.0.0)^98^ command bamCompare –operation mean with default parameters.

Peak-calling was done with MACS2 (v2.1.1.20160309)^100^ on individual replicate with the default parameters. For each genotype, we then generated a genotype specific high confidence peakset by intersection of peaks called in two replicates. IDR framework (v2.0.4)^101^ were used to generate quality metrics for the processed ChIPseq data. R package DiffBind (v3.0.15)^102,103^ and DEseq2 (v1.30.1)^104^ were then used to re-center peaks to ±200bp regions surrounding summit, generate normalized binding intensity matrix, and differential binding matrix. We defined differentially bound peaks between each mutant and wild type sample if the absolute log_2_FC is more than 1.0, corresponding to two-fold, and the FDR is smaller than 5e-2.

To associate peaks to genes, we used Genomic Regions Enrichment of Annotations Tool (GREAT v4.0.4)^105^ through the R package rGREAT (v1.19.2)^106^. Each peak was assigned to the closest TSS within 100kb.

#### Binding intensity heatmap and clustering

To generate the binding intensity heatmap in Figure 2A, we first compiled the genotype specific high confidence peakset for all genotypes into a single consensus peakset and only peaks with at least 5 cpm in all genotypes were retained. Python package fastcluster (v1.1.26)^107^ was used to perform hierarchical clustering of the consensus peakset intensity matrix with parameters method=’single’, metric=’euclidean’. The genomic regions corresponding to the two major clusters were exported and used to generate binding intensity heatmaps with deeptools (v3.0.0)^98^.

#### Genomic region enrichment of CRX peaks

Peak annotation in Figure 2D were obtained using annotatePeaks.pl from HOMER (v4.8).

#### De novo motif searching

The mm10 fasta sequences for each genotype specific peaks were obtained using R package BSgenome (v 1.58.0)^108^. De novo motif enrichment analysis for each set of sequences were then performed with MEME-ChIP in MEME Suite (v5.0.4)^109^ using order 1 Markov background model and default parameters. Since homeodomain motifs are relatively short and can be repetitive (e.g. K88N motif), we reported DREME^38^ found motifs for Figure 2E, which is more sensitive than MEME to find short, repetitive motifs.

#### RNA-seq data analysis

2×150bp reads from Illumina NovaSeq were obtained for all samples with a minimum depth of 17 M reads at Novogene (Beijing, China). Sequencing reads were first run through Trim Galore (v0.6.1)^92^ to remove adapter sequences and then QC by FastQC (v0.11.5)^93^. Trimmed reads were then mapped to the mm10 genome and quantified with kallisto (v0.46.2)^110^. Kallisto output transcript-level abundance matrices were then imported and summarized into gene-level matrices with R package tximport (v1.18.0)^111^. DEseq2 (v1.30.1)^104^ was then used for normalization and differential expression analysis. The normalized count and differential expression matrices were then exported and further processed with customized python scripts.

We defined differentially expressed genes between each mutant and wild type sample if the absolute log_2_FC is more than 1.0, corresponding to two-fold, and the FDR is smaller than 1e-2. For comparison between heterozygous and homozygous mutants, we first filtered genes with at least 5 cpm and then those that were called differentially expressed compared with wild type in at least one mutant genotype. We retrieved gene names in Supplementary Tables S3-S5 from the Database for Annotation, Visualization and Integrated Discovery (DAVID, v6.8)^112^.

#### Definition of CRX-dependent and -independent gene set

We first identified CRX peaks that were bound in the *WT* sample but lost in *R90W/W* sample (log_2_FC < -1 and FDR < 5e-2). This yielded a total of 7677 peaks. We then found the genes associated with these peaks. We defined a gene to be CRX-dependent activated if its expression was down in adult (P21) *R90W/W* RNA-seq sample (log_2_FC < -0.6 and FDR < 1e-5). Similarly, a gene is defined as CRX-dependent suppressed if its expression was up in adult *R90W/W* RNA-seq sample (log_2_FC > 0.6 and FDR < 1e-5). A gene is defined as CRX-independent if its expression was not significantly affected in adult *R90W/W* RNA-seq samples. There were 617 CRX-dependent activated, 135 CRX-dependent suppressed, and 5565 CRX-independent genes. Manual inspection of the CRX-dependent suppressed genes revealed no clear association with photoreceptor development. Therefore, we did not further pursue this gene set. The complete list of CRX-dependent activated genes that showed differential expression in at least one of the HD mutant retinas can be found in Supplementary Table S5. The lists for CRX-independent genes that showed differential expression in *Crx^E80A^*or *Crx^K88N^* mutant retinas can be found in Supplementary Tables S6 and S7 respectively.

#### Gene ontology analysis

Gene ontology analysis in Figures S4B, S4E, and S5F were performed using R package clusterProfiler (v4.0.5)^113,114^ with the genome wide annotation package org.Mm.eg.db (v3.12.0)^115^. Redundant enriched GO terms were removed using simplify() function with parameters cutoff=0.7, by=“p.adjust”. The enrichment analysis results were then exported in table format and further processed for plotting with python.

#### Aldiri et al. RNA-seq data re-analysis

The RNA-seq data from *Aldiri* et al.^39^ were obtained from GEO under accession numbers GSE87064. The reads were processed similarly as all other RNA-seq data generated in this study. For Figures S4C, S4F, S5C, S5E,S6A, S6B, expression row z-scores were calculated using average cpm from replicates at each age.

#### Statistical analysis

One-way ANOVA with Turkey honestly significant difference (HSD) test in Figures 6A, S3C, and S7D were performed with python packages scipy (v1.8.1)^116^ and scikit_posthocs (v0.7.0)^117^. Two-sided Mann-Whitney U test in Figure S5A was performed with python package scipy (v1.8.1)^116^.

## Key resources table

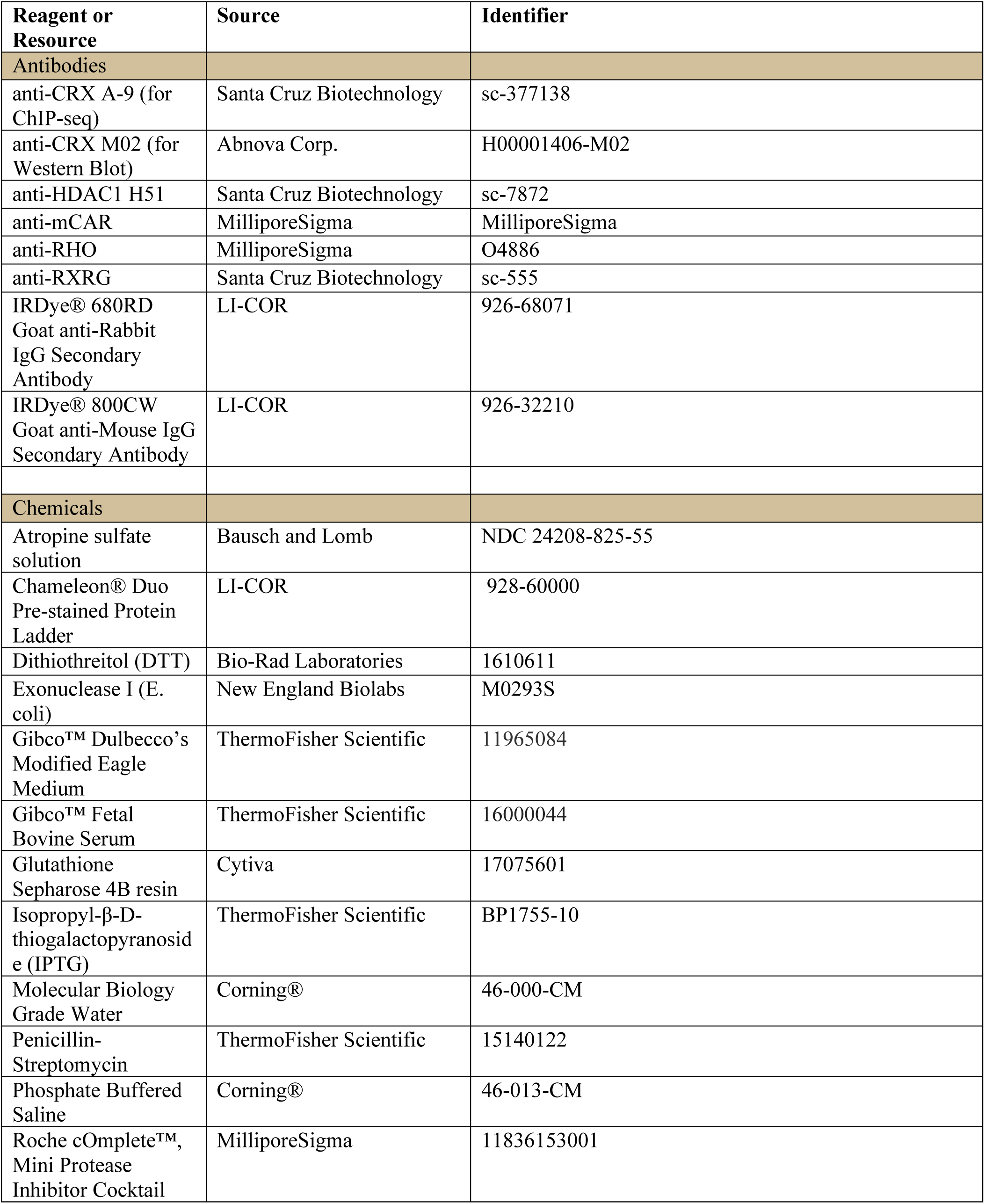
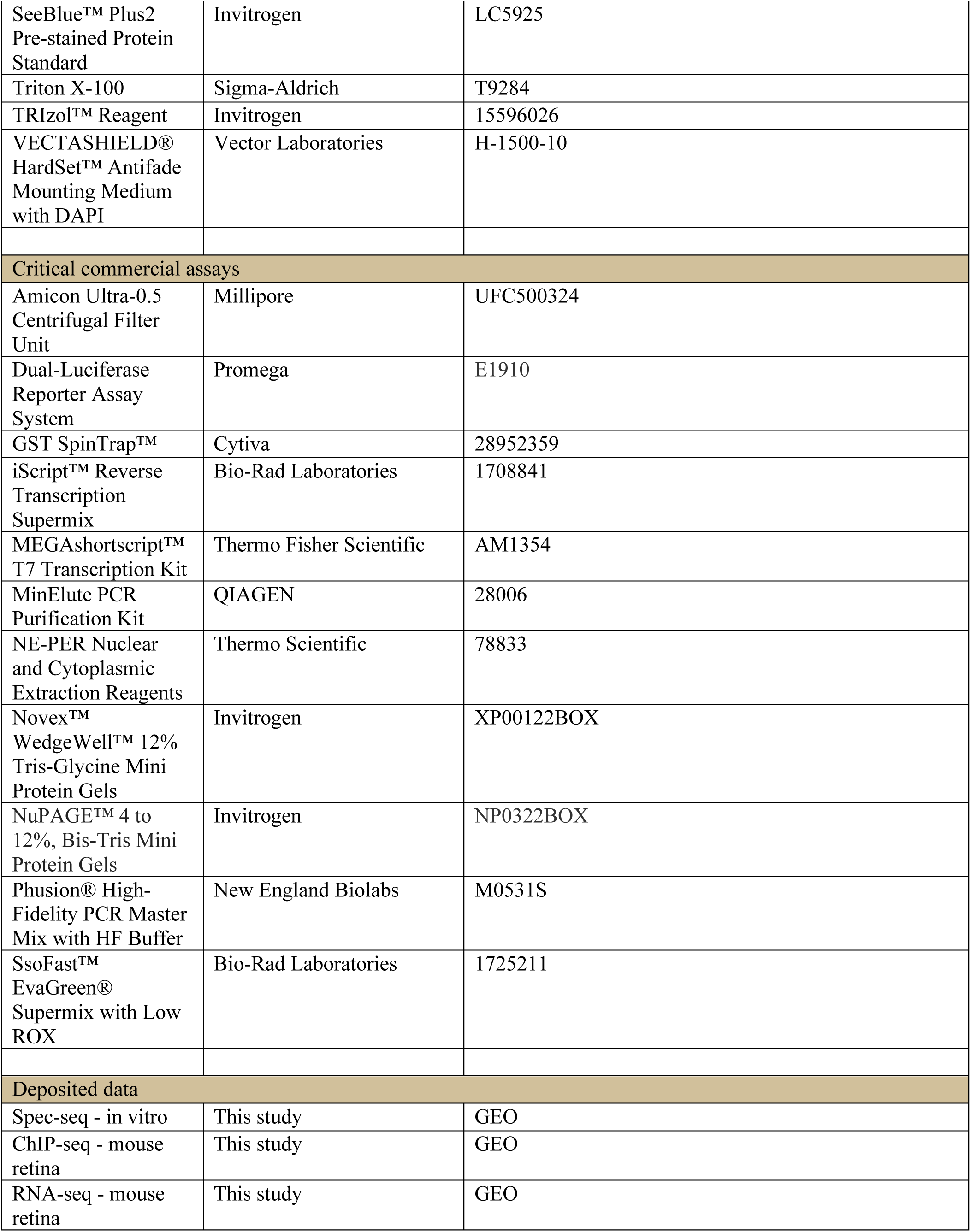
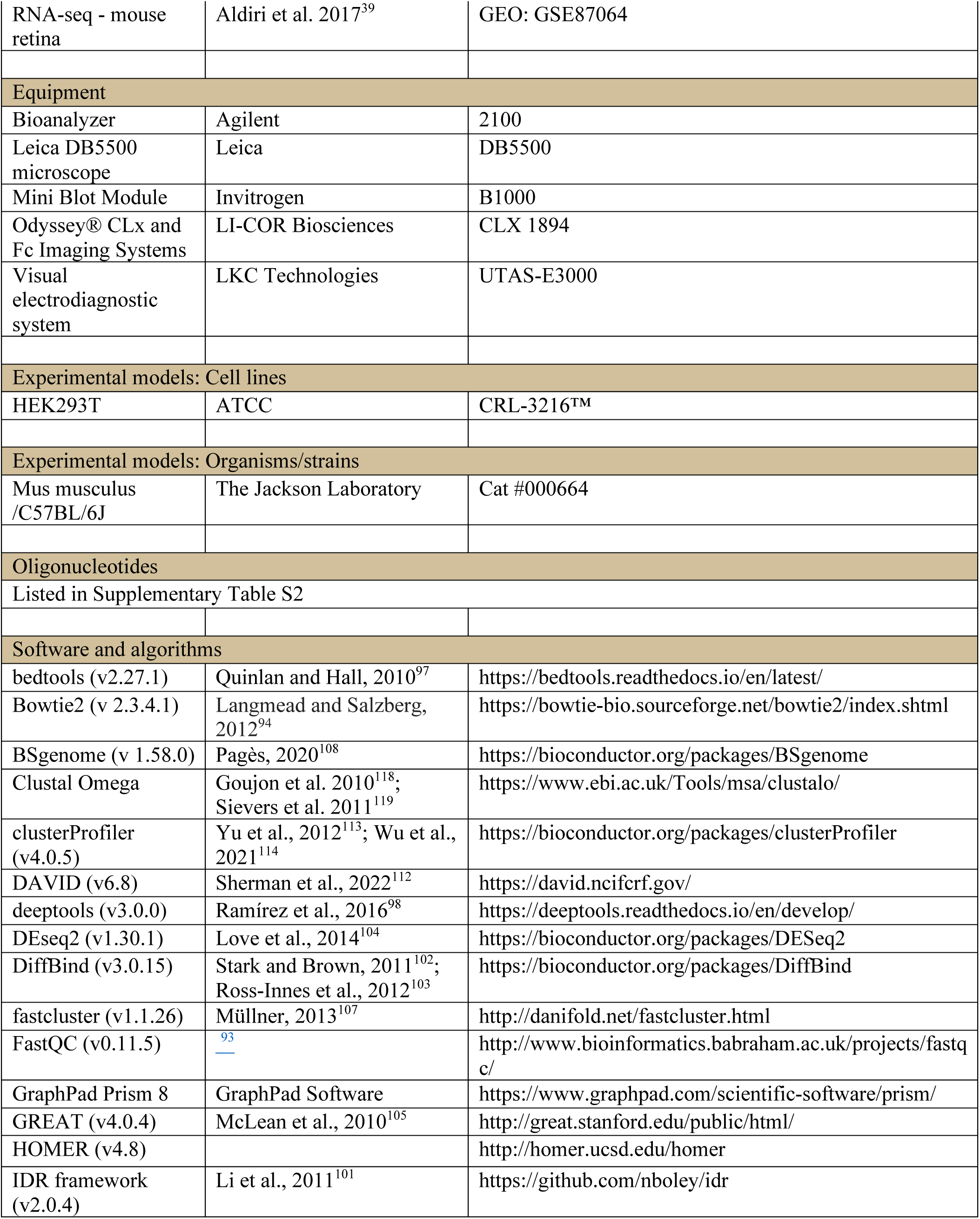
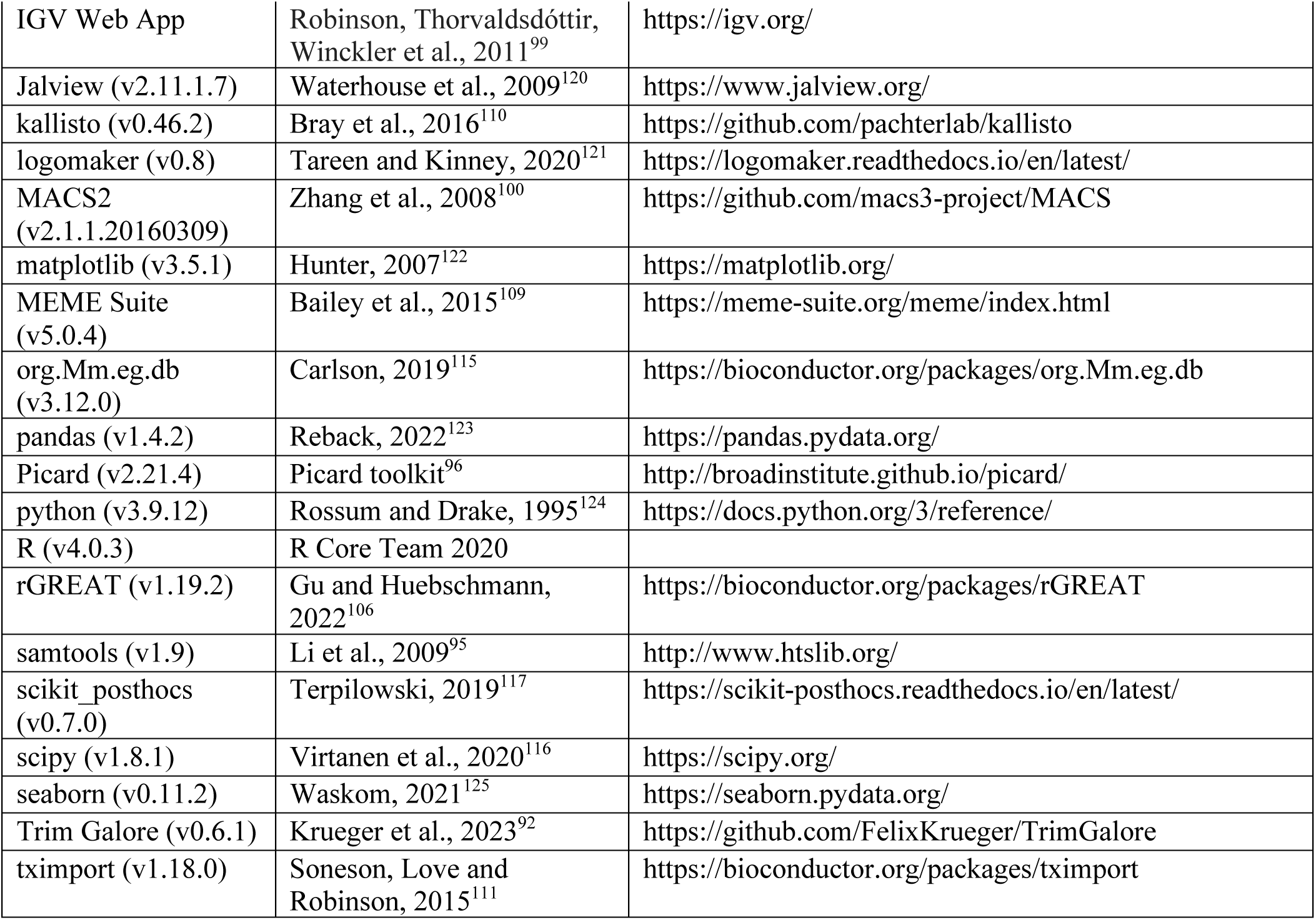

## Supplementary tables

Supplementary Table S1. HD TF accession numbers (related to Figure 1B)

Supplementary Table S2. Plasmids and primers for biochemistry experiments

Supplementary Table S3. Spec-seq ePWMs (related to Figures 1G, 1K, 1L, 1M, and S2I-S2L)

Supplementary Table S4. DNA sequences for mutation knock-in mice generation

Supplementary Table S5. CRX-dependent activated genes differentially expressed in at least one mutant mouse model (ordered as in Figure 3B)

Supplementary Table S6. CRX-independent genes mis-regulated in *Crx^E80A^* mutants (ordered as in Figure S5C)

Supplementary Table S7. CRX-independent genes mis-regulated in *Crx^K88N^* mutants (ordered as in Figure S5E)

Supplementary Table S8. Annotation for phototransduction genes in Figure 4B

Supplementary Table S9 Quantification and statistical analysis (related to Figures S3C and S5A)

---

**Supplementary Figure S1.**
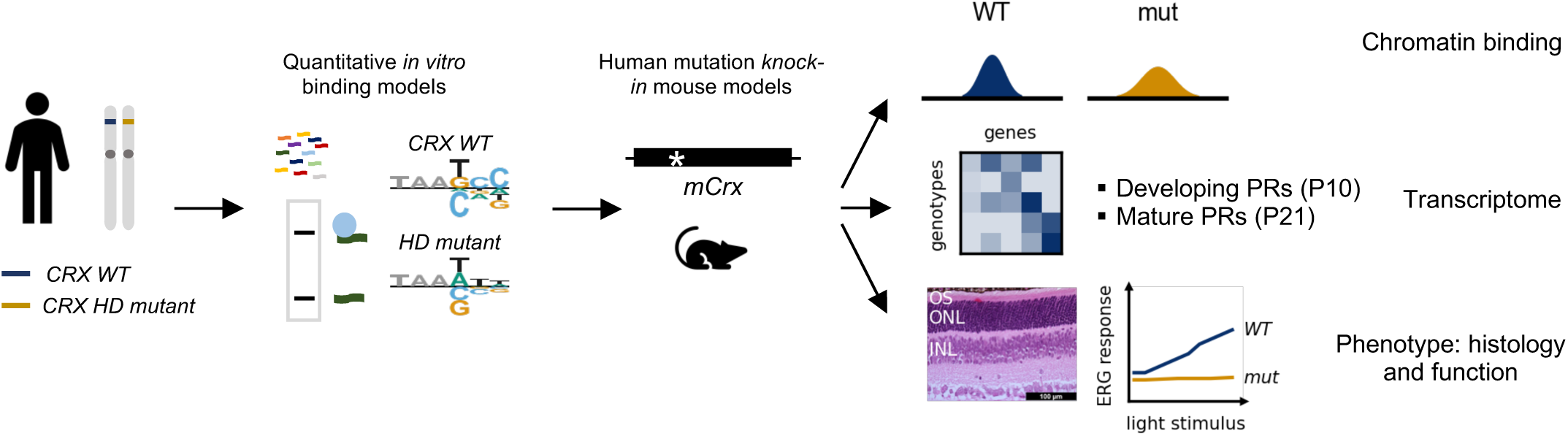
Multi-omics approach to investigate the functional consequences of dominant disease mutations on CRX regulatory activities and photoreceptor development. Human retinopathy associated CRX HD mutant is first tested *in vitro* for HD-DNA interactions by Spec-seq. Quantitative binding models are generated for WT and mutant HDs. Each mutation is then introduced into endogenous *mCrx* locus to generate human mutation knock-in mouse models. ChIP-seq is employed to characterize CRX chromatin binding in *WT* and mutant mouse retinas. Bulk RNA-seq is then applied to determine transcriptomic changes in developing (P10) and mature (P21) photoreceptors (PRs) in both *WT* and mutant mouse retinas. Last, phenotypic characterization on retinal morphology and visual functions is carried out to understand the consequences of mutant CRX chromatin binding and associated transcriptomic alterations.

**Supplementary Figure S2.**
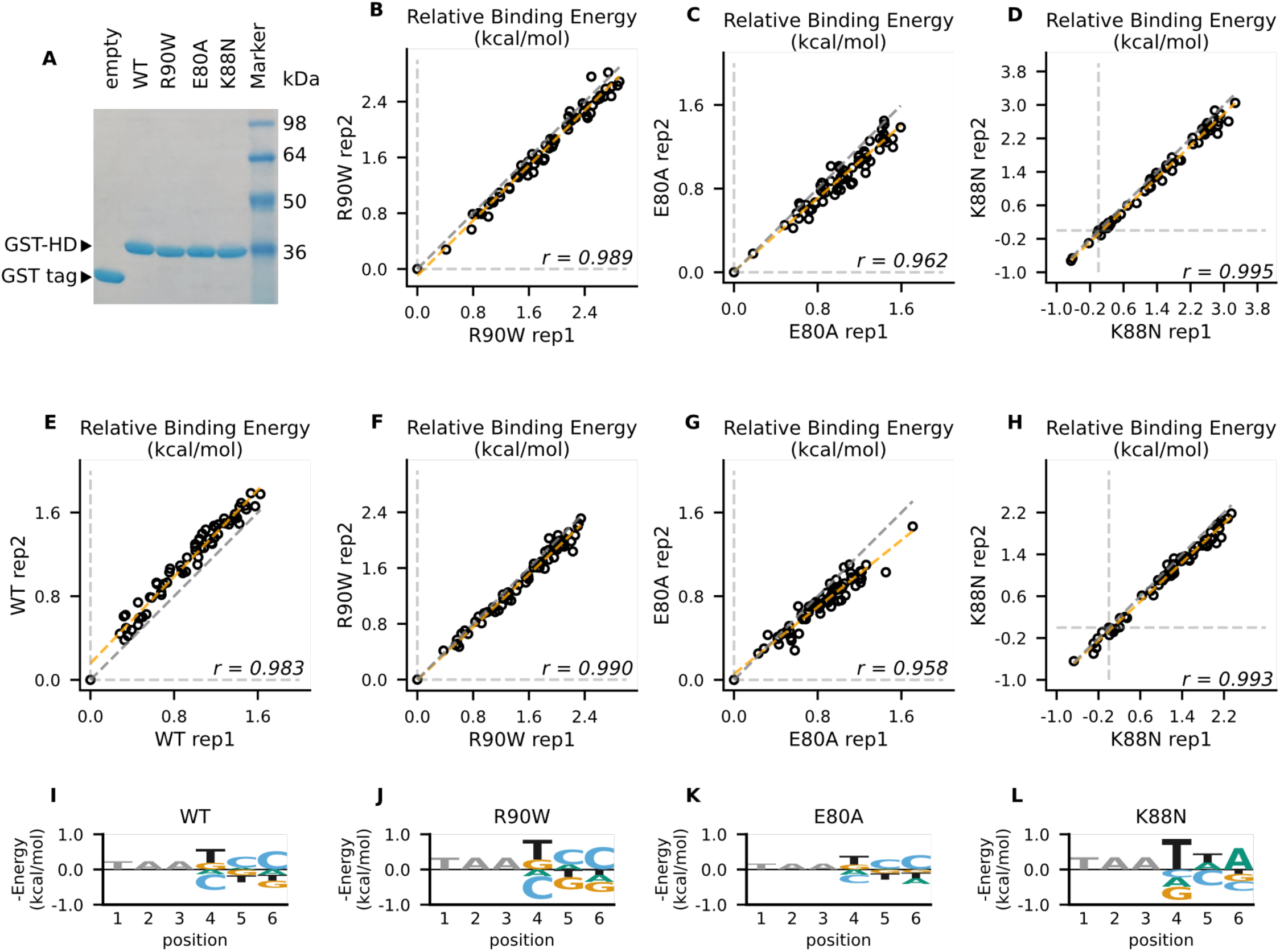
Reversed-strand Spec-seq library showed similar changes in mutant CRX HD DNA-binding specificity. (A) Native SDS-PAGE gel image of affinity purified empty GST tag and GST-CRX HDs. (B-D) Relative binding energy comparison from two different experiments for R90W HD (B), E80A HD (C), and K88N HD (D) on the same Spec-seq library as in Figure 1. (E-L) Spec-seq experiments of a second library with the TAANNN sites on the reverse strand show similar results. (E-H) Relative binding energy comparison from two different experiments for WT HD (E), R90W HD (F), E80A HD (G), and K88N HD (H) on the reversed monomeric library. The identity line is represented in grey dash. The orange dashed line shows the best linear fit to the data. (I-L) Binding energy models for WT HD (I), R90W HD (J), E80A HD (K), and K88N HD (L) obtained from the reversed monomeric library. Quantitative difference with models obtained from forward-oriented library likely comes from difference in sequences immediately flanking the TAANNN variable region.

**Supplementary Figure S3.**
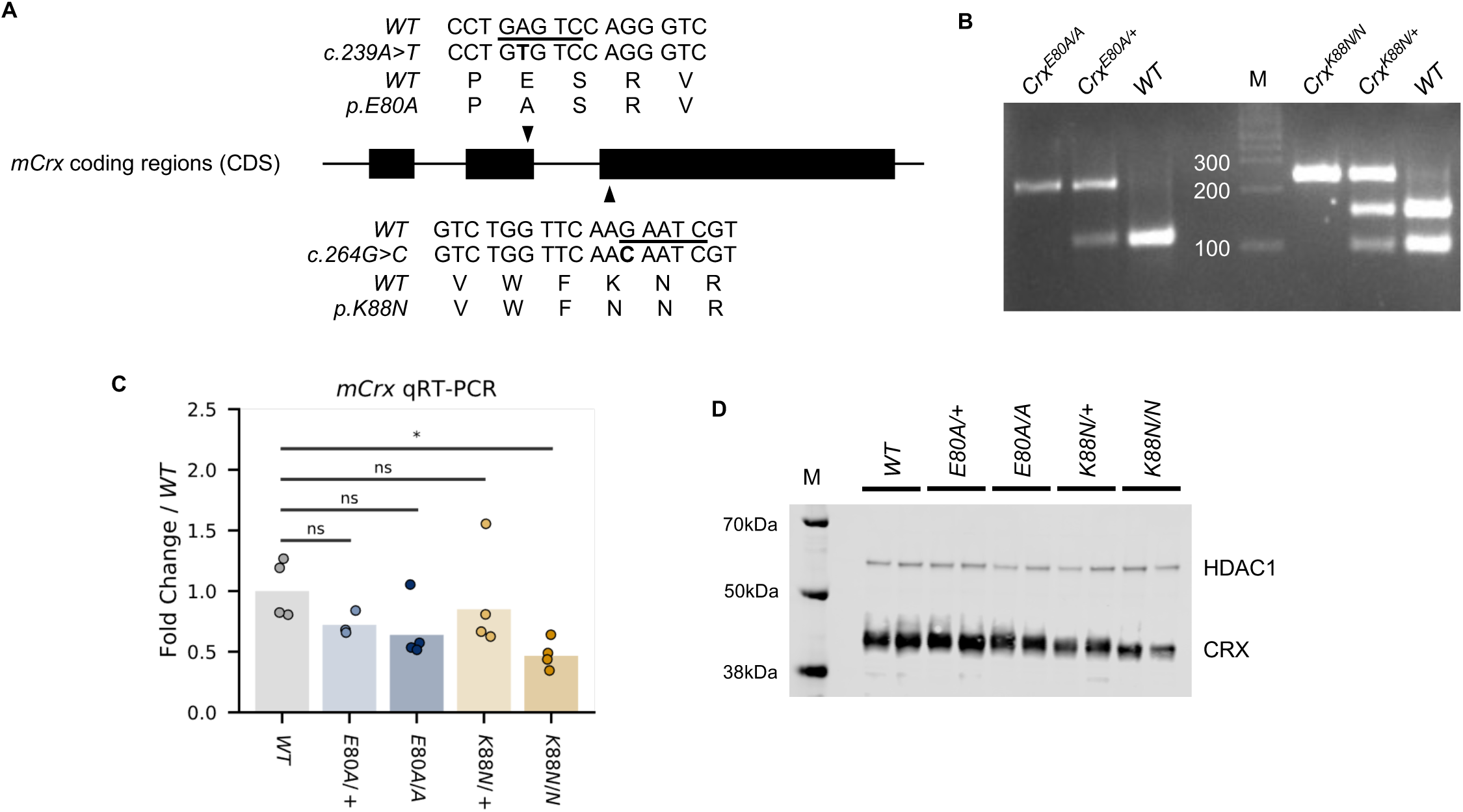
WT and mutation knock-in mouse CRX sequences and genotyping identifications. (A) Alignment of *mCrx* cDNA and protein sequences showing the nucleotide substitutions and amino acid changes of *Crx^E80A^* (top) and *Crx^K88N^* (bottom) alleles. Only coding regions of *mCrx* exons are shown and the diagram is not to scale. Underlined bases in WT sequences indicate the restriction enzyme HinfI cut sites used in the genotyping PCR. (B) Representative mutation knock-in mouse genotyping gel image. (C) Barchart and stripplot showing *mCrx* mRNA expression levels in P14 *WT* and mutant mouse retinas. *P-values* for one-way ANOVA with Turkey honestly significant difference (HSD) test are indicated. *p-value*: ****: ≤0.0001, ***: ≤0.001, **: ≤0.01, *: ≤0.05, ns: >0.05. (D) Immunoblots of nuclear extracts obtained from P14 *WT* and mutant mouse retinas showing that full-length CRX protein are produced and localized to the nucleus fraction in all mutant mouse retinas. HDAC1 was used as a loading control.

**Supplementary Figure S4.**
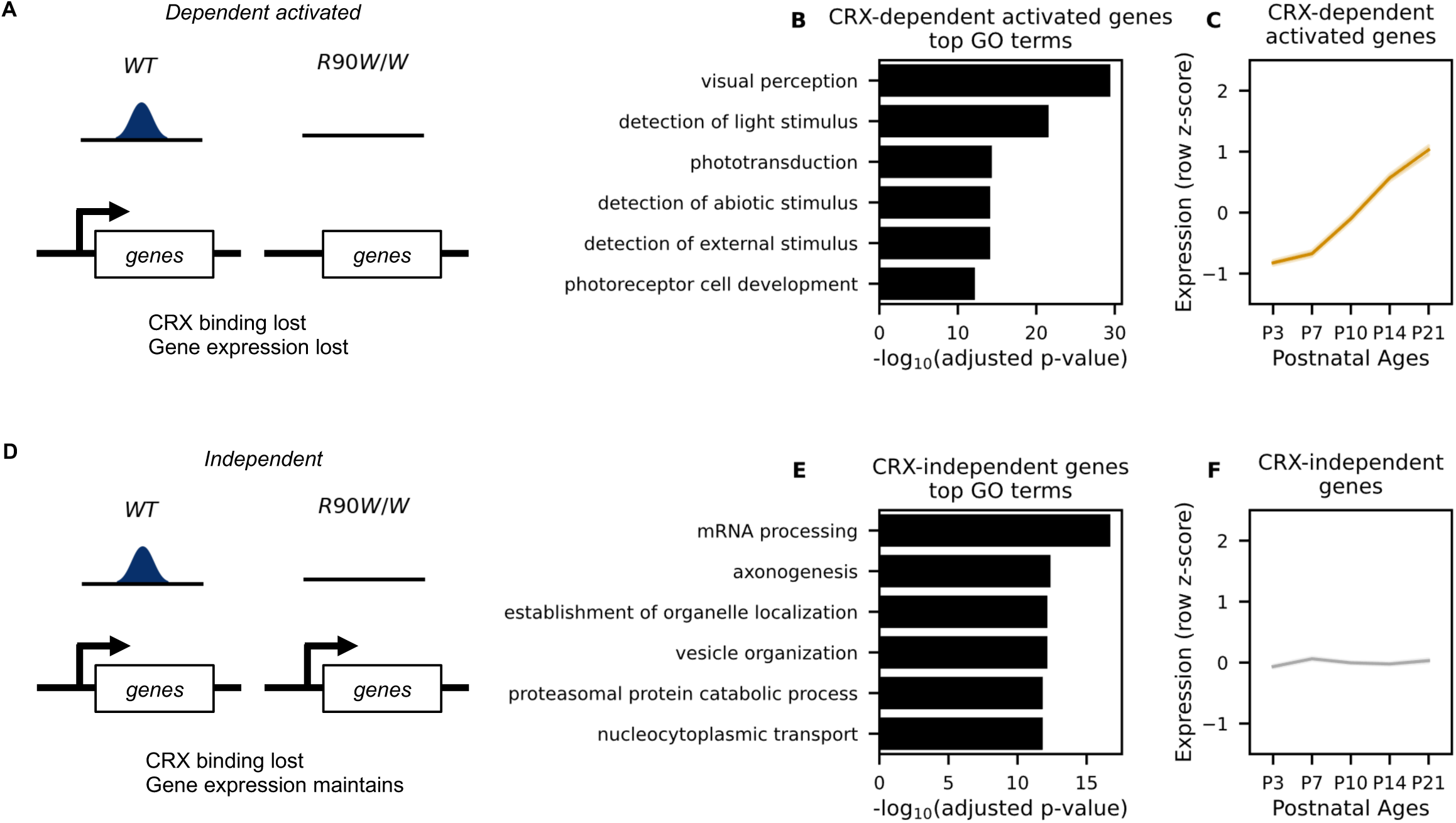
Definition of CRX-dependent activated and CRX-independent gene sets. (A) Schematic representation of CRX-dependent activated genes where CRX binding nearby is required for the expression of these genes in mature *WT* retinas. (B) Top GO terms associated with CRX-dependent activated genes. Benjamini-Hochberg adjusted p-values are shown. (C) Line plot showing average expression pattern of CRX-dependent activated genes during normal post-natal retina. (D) Definition of CRX-independent genes where CRX binding nearby is dispensable for the expression of these genes in mature *WT* retinas. (E) Top GO terms associated with CRX-independent genes. (F) Line plot showing average expression pattern of CRX-independent genes during normal retina development. RNA-seq data in (C) and (F) were retrieved from Aldiri et al.^39^ (GEO accession: GSE87064).

**Supplementary Figure S5.**
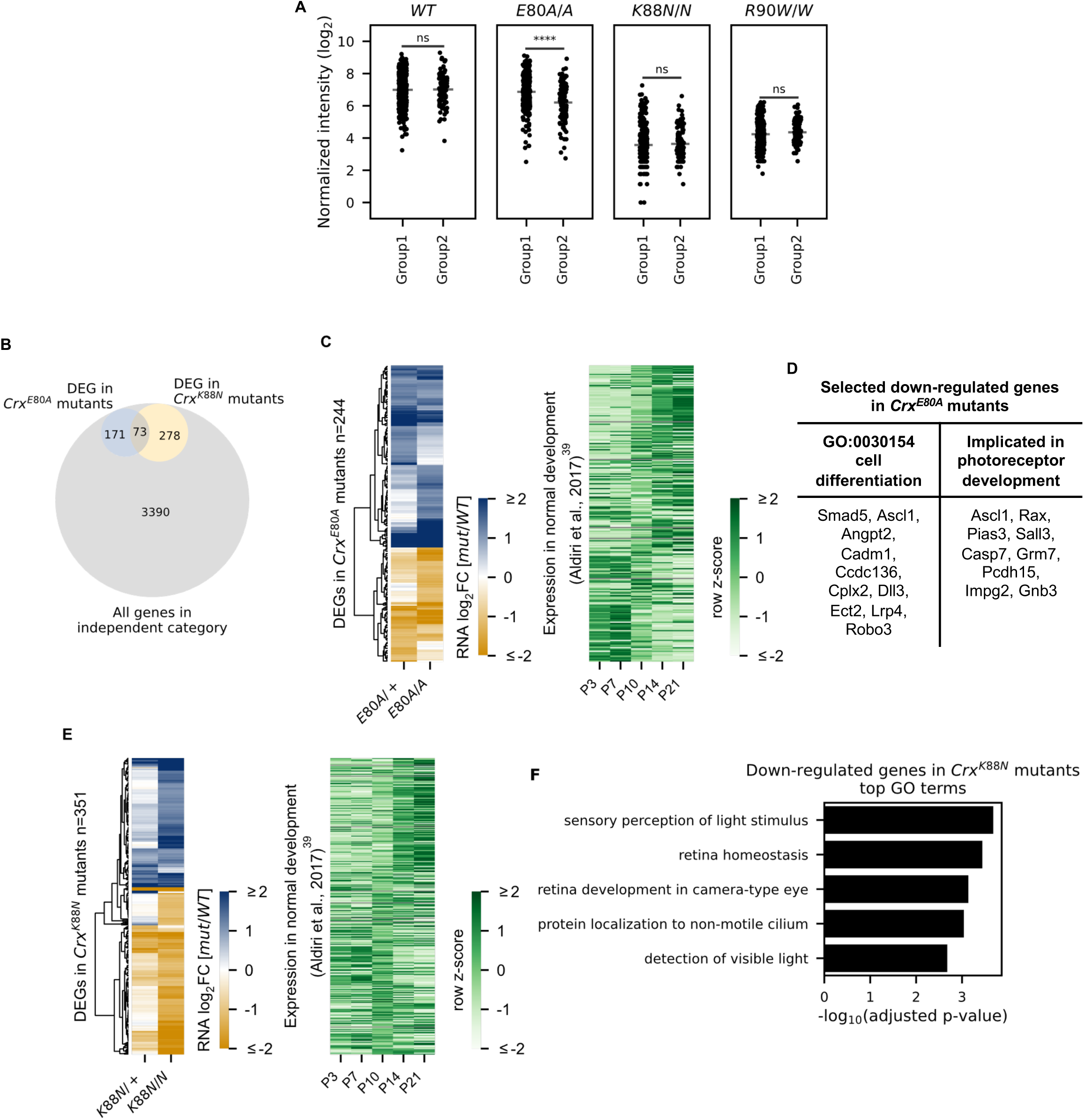
E80A and K88N mutation each causes novel gene expression changes in the CRX-independent category. (A) Strip plots showing normalized CRX ChIP-seq intensity at peaks associated with Group1 or Group2 genes in WT and CRX mutant mouse retinas. *P-values* for two-sided Mann-Whitney U test are indicated. *p-value*: ****: ≤0.0001, ***: ≤0.001, **: ≤0.01, *: ≤0.05, ns: >0.05. (B) Venn diagram showing the overlap of genes differentially expressed (DEGs) in *Crx^E80A^* (pale blue) and *Crx^K88N^* (pale yellow) but not in *Crx^R90W/W^* (grey) mutant retinas. For DEGs in *Crx^E80A^* and *Crx^K88N^* mutants, genes that were differentially expressed in either heterozygotes or homozygotes, or both were counted. (C) Heat map showing the expression changes of CRX-independent genes that are DEGs in at least one of the *Crx^E80A^* mutants (n = 244, left). Heat map on the right shows the expression pattern of these genes during normal post-natal development (data from GSE87064). (D) Table showing selected genes down-regulated in *Crx^E80A^* mutants that have been implicated in cell differentiation or photoreceptor development. (E) Heat map showing the expression changes of CRX-independent genes that are DEGs in at least one of the *Crx^K88N^* mutants (n = 351, left). Heat map on the right shows the expression pattern of these genes during normal post-natal development (data from GSE87064). (F) Bar chart showing GO term enrichment of DEGs in *Crx^K88N^* mutants.

**Supplementary Figure S6.**
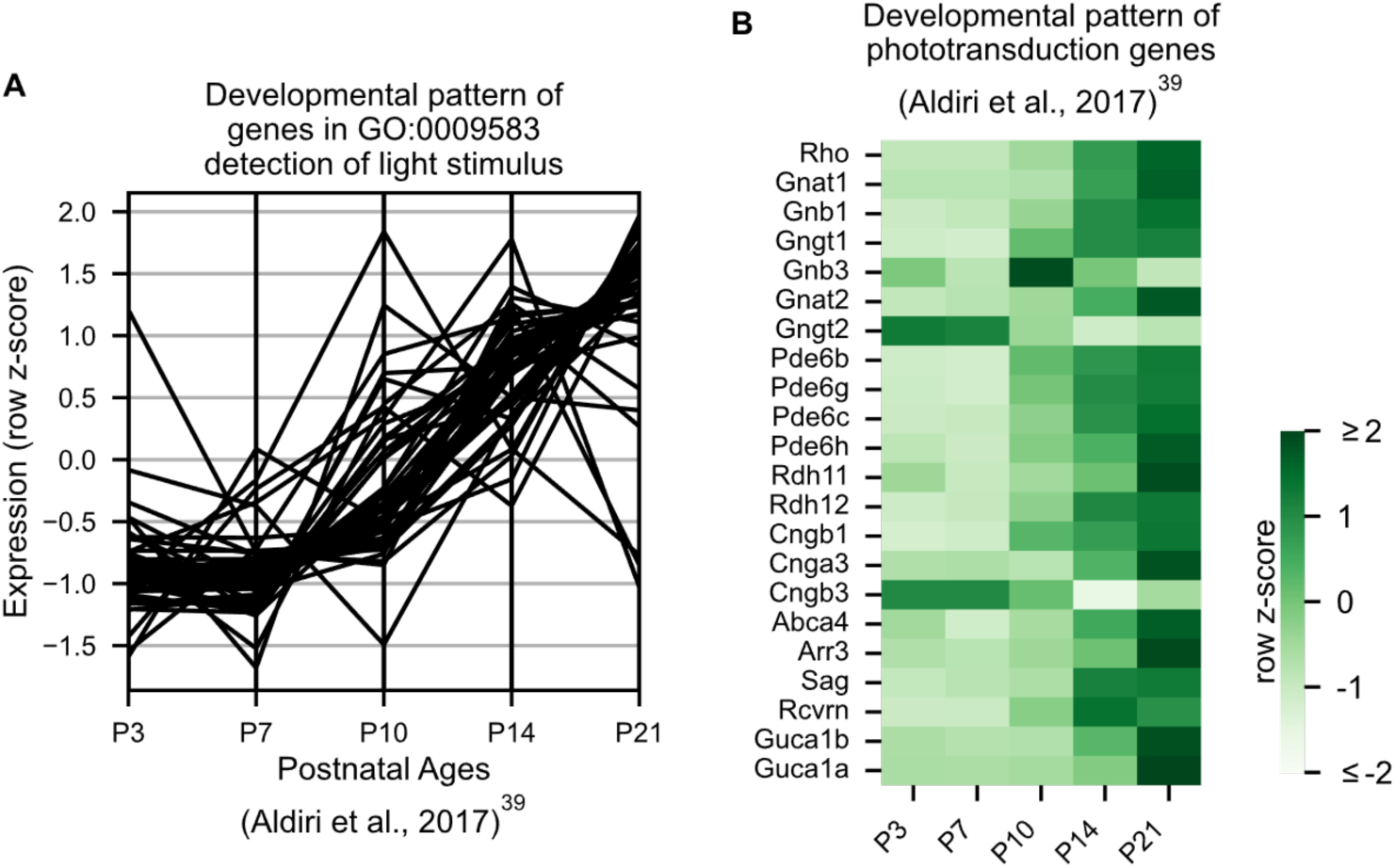
Developmental expression pattern of phototransduction genes in *WT* animals. (A) Parallel coordinates plot showing expression pattern of genes in GO:0009583 during normal post-natal retina development (data from GSE87064). Row z-score is shown. (B) Heatmap showing expression patterns of phototransduction genes during normal post-natal retina development (data from GSE87064).Row z-score is shown.

**Supplementary Figure S7.**
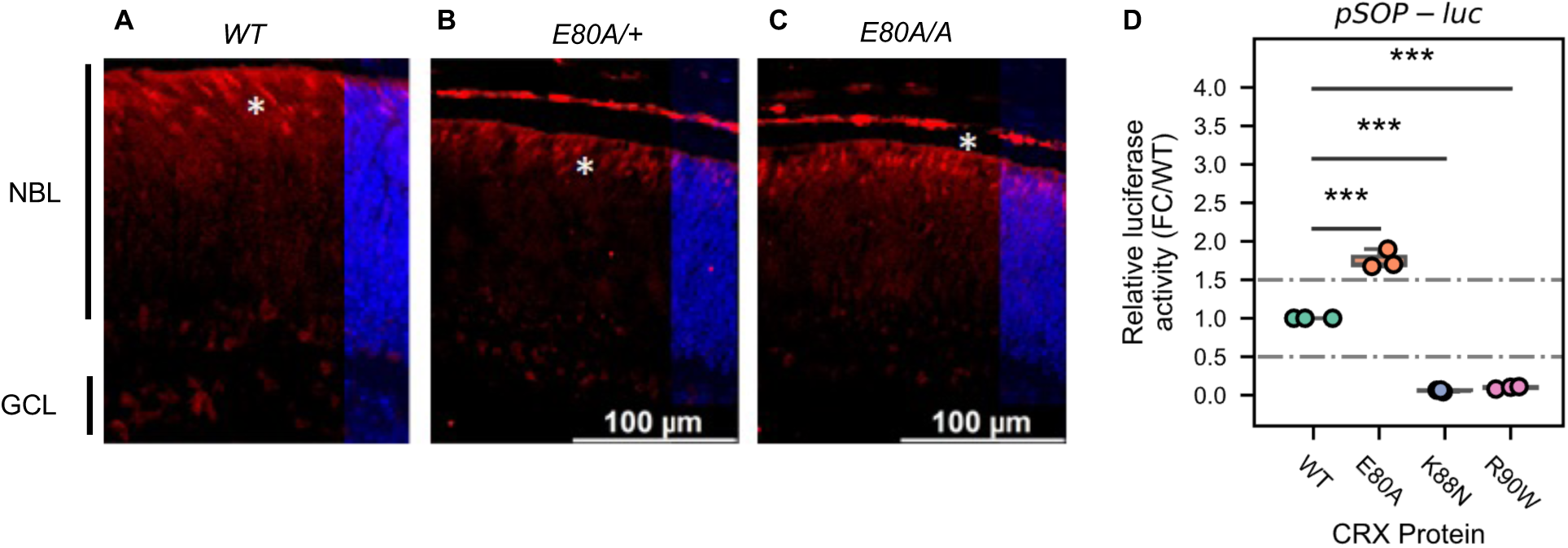
Cone photoreceptors born in *Crx^E80A^* retinas and hyperactivity of CRX E80A at S-opsin promoter. (A-C) Immunostaining shows that Retinoid X receptor gamma (RXR*γ*, red), a fated cone photoreceptor marker, is present in P0 *WT, Crx^E80A/+^* and *Crx^E80A/A^* retinas. Nuclei are visualized by DAPI staining (Blue). Asterisks indicate examples of RXRG+ cells. NBL: neuroblast layer; GCL: ganglion cell layer. Scale bar, 100µm. (D) Boxplot showing luciferase reporter activities of different CRX variants at the *S-opsin* promoter sequences. *P-values* for one-way ANOVA with Turkey honestly significant difference (HSD) test are indicated. *p-value*: ****: ≤0.0001, ***: ≤0.001, ns: >0.05.

## Reference

1. Mark, M., Rijli, F.M. & Chambon, P. Homeobox Genes in Embryogenesis and Pathogenesis. Pediatric Research 42, 421–429 (1997).

2. Lewis, E.B. A gene complex controlling segmentation in Drosophila. Nature 276, 565–570 (1978).

3. Leung, R.F. et al. Genetic Regulation of Vertebrate Forebrain Development by Homeobox Genes. Frontiers in Neuroscience 16(2022).

4. Chi, Y.-I. Homeodomain revisited: a lesson from disease-causing mutations. Human Genetics 116, 433–444 (2005).

5. Zagozewski, J.L., Zhang, Q., Pinto, V.I., Wigle, J.T. & Eisenstat, D.D. The role of homeobox genes in retinal development and disease. Developmental Biology 393, 195–208 (2014).

6. Diacou, R. et al. Cell fate decisions, transcription factors and signaling during early retinal development. Progress in Retinal and Eye Research 91, 101093 (2022).

7. Beby, F. & Lamonerie, T. The homeobox gene Otx2 in development and disease. Experimental Eye Research 111, 9–16 (2013).

8. Henderson, R.H. et al. A rare de novo nonsense mutation in OTX2 causes early onset retinal dystrophy and pituitary dysfunction. Mol Vis 15, 2442–7 (2009).

9. Abouzeid, H. et al. PAX6 aniridia and interhemispheric brain anomalies. Mol Vis 15, 2074–83 (2009).

10. Lima Cunha, D., Arno, G., Corton, M. & Moosajee, M. The Spectrum of PAX6 Mutations and Genotype-Phenotype Correlations in the Eye. Genes (Basel) 10(2019).

11. Voronina, V.A. et al. Mutations in the human RAX homeobox gene in a patient with anophthalmia and sclerocornea. Human Molecular Genetics 13, 315–322 (2003).

12. Abouzeid, H. et al. RAX and anophthalmia in humans: evidence of brain anomalies. Mol Vis 18, 1449–56 (2012).

13. Muranishi, Y. et al. An essential role for RAX homeoprotein and NOTCH-HES signaling in Otx2 expression in embryonic retinal photoreceptor cell fate determination. J Neurosci 31, 16792–807 (2011).

14. Chen, S. et al. Crx, a Novel Otx-like Paired-Homeodomain Protein, Binds to and Transactivates Photoreceptor Cell-Specific Genes. Neuron 19, 1017–1030 (1997).

15. Furukawa, T., Morrow, E.M. & Cepko, C.L. Crx, a Novel otx-like Homeobox Gene, Shows Photoreceptor-Specific Expression and Regulates Photoreceptor Differentiation. Cell 91, 531–541 (1997).

16. Furukawa, T., Morrow, E.M., Li, T., Davis, F.C. & Cepko, C.L. Retinopathy and attenuated circadian entrainment in Crx-deficient mice. Nature Genetics 23, 466 (1999).

17. Chau, K.Y., Chen, S., Zack, D.J. & Ono, S.J. Functional domains of the cone-rod homeobox (CRX) transcription factor. J Biol Chem 275, 37264–70 (2000).

18. Chen, S. et al. Functional analysis of cone–rod homeobox (CRX) mutations associated with retinal dystrophy. Human Molecular Genetics 11, 873–884 (2002).

19. Tran, N.M. et al. Mechanistically Distinct Mouse Models for CRX-Associated Retinopathy. PLOS Genetics 10, e1004111 (2014).

20. Ruzycki, P.A., Tran, N.M., Kolesnikov, A.V., Kefalov, V.J. & Chen, S. Graded gene expression changes determine phenotype severity in mouse models of CRX-associated retinopathies. Genome Biology 16, 171 (2015).

21. Freund, C.L. et al. Cone-Rod Dystrophy Due to Mutations in a Novel Photoreceptor-Specific Homeobox Gene (CRX) Essential for Maintenance of the Photoreceptor. Cell 91, 543–553 (1997).

22. Swaroop, A. et al. Leber Congenital Amaurosis Caused by a Homozygous Mutation (R90W) in the Homeodomain of the Retinal Transcription Factor CRX: Direct Evidence for the Involvement of CRX in the Development of Photoreceptor Function. Human Molecular Genetics 8, 299–305 (1999).

23. Nichols, L.L., 2nd et al. Two novel CRX mutant proteins causing autosomal dominant Leber congenital amaurosis interact differently with NRL. Hum Mutat 31, E1472–83 (2010).

24. Desplan, C., Theis, J. & O’Farrell, P.H. The sequence specificity of homeodomain-DNA interaction. Cell 54, 1081–1090 (1988).

25. Trelsman, J., Gönczy, P., Vashishtha, M., Harris, E. & Desplan, C. A single amino acid can determine the DNA binding specificity of homeodomain proteins. Cell 59, 553–562 (1989).

26. Hanes, S.D. & Brent, R. DNA specificity of the bicoid activator protein is determined by homeodomain recognition helix residue 9. Cell 57, 1275–1283 (1989).

27. Hanes, S.D. & Brent, R. A genetic model for interaction of the homeodomain recognition helix with DNA. Science 251, 426 (1991).

28. Ades, S.E. & Sauer, R.T. Specificity of Minor-Groove and Major-Groove Interactions in a Homeodomain-DNA Complex. Biochemistry 34, 14601–14608 (1995).

29. Wilson, D.S., Guenther, B., Desplan, C. & Kuriyan, J. High resolution crystal structure of a paired (Pax) class cooperative homeodomain dimer on DNA. Cell 82, 709–719 (1995).

30. Gehring, W.J., et al. Homeodomain-DNA recognition. Cell 78, 211–223 (1994).

31. Noyes, M.B. et al. Analysis of homeodomain specificities allows the family-wide prediction of preferred recognition sites. Cell 133, 1277–89 (2008).

32. Stormo, G.D., Zuo, Z. & Chang, Y.K. Spec-seq: determining protein–DNA-binding specificity by sequencing. Briefings in Functional Genomics 14, 30–38 (2015).

33. Zuo, Z., Roy, B., Chang, Y.K., Granas, D. & Stormo, G.D. Measuring quantitative effects of methylation on transcription factor-DNA binding affinity. Science advances 3, eaao1799–eaao1799 (2017).

34. Zuo, Z. & Stormo, G.D. High-Resolution Specificity from DNA Sequencing Highlights Alternative Modes of Lac Repressor Binding. Genetics 198, 1329–1343 (2014).

35. Corbo, J.C. et al. CRX ChIP-seq reveals the cis-regulatory architecture of mouse photoreceptors. Genome Res 20, 1512–25 (2010).

36. Kwasnieski, J.C., Mogno, I., Myers, C.A., Corbo, J.C. & Cohen, B.A. Complex effects of nucleotide variants in a mammalian cis;-regulatory element. Proceedings of the National Academy of Sciences 109, 19498 (2012).

37. Bassett, E.A. & Wallace, V.A. Cell fate determination in the vertebrate retina. Trends Neurosci 35, 565–73 (2012).

38. Bailey, T.L. DREME: motif discovery in transcription factor ChIP-seq data. Bioinformatics 27, 1653–1659 (2011).

39. Aldiri, I. et al. The Dynamic Epigenetic Landscape of the Retina During Development, Reprogramming, and Tumorigenesis. Neuron 94, 550–568 e10 (2017).

40. Kim, J.W. et al. NRL-Regulated Transcriptome Dynamics of Developing Rod Photoreceptors. Cell Rep 17, 2460–2473 (2016).

41. Onishi, A. et al. The orphan nuclear hormone receptor *ERRβ* controls rod photoreceptor survival. Proceedings of the National Academy of Sciences 107, 11579 (2010).

42. Yoshida, S. et al. Expression profiling of the developing and mature Nrl-/- mouse retina: identification of retinal disease candidates and transcriptional regulatory targets of Nrl. Hum Mol Genet 13, 1487–503 (2004).

43. Zang, J. & Neuhauss, S.C.F. The Binding Properties and Physiological Functions of Recoverin. Frontiers in Molecular Neuroscience 11(2018).

44. Palczewski, K. G protein-coupled receptor rhodopsin. Annu Rev Biochem 75, 743–67 (2006).

45. Dryja, T.P., Hahn, L.B., Reboul, T. & Arnaud, B. Missense mutation in the gene encoding the α subunit of rod transducin in the Nougaret form of congenital stationary night blindness. Nature Genetics 13, 358–360 (1996).

46. Carrigan, M. et al. A novel homozygous truncating *GNAT1* mutation implicated in retinal degeneration. British Journal of Ophthalmology 100, 495 (2016).

47. Dvir, L. et al. Autosomal-Recessive Early-Onset Retinitis Pigmentosa Caused by a Mutation in *PDE6G*, the Gene Encoding the Gamma Subunit of Rod cGMP Phosphodiesterase. The American Journal of Human Genetics 87, 258–264 (2010).

48. Allikmets, R. et al. Mutation of the Stargardt Disease Gene (ABCR) in Age-Related Macular Degeneration. Science 277, 1805–1807 (1997).

49. Nasonkin, I. et al. Mapping of the rod photoreceptor ABC transporter (ABCR) to 1p21–p22.1 and identification of novel mutations in Stargardt’s disease. Human Genetics 102, 21–26 (1998).

50. Kitamura, E. et al. Disruption of the Gene Encoding the β1-Subunit of Transducin in the Rd4/+ Mouse. Investigative Ophthalmology & Visual Science 47, 1293–1301 (2006).

51. Janecke, A.R. et al. Mutations in RDH12 encoding a photoreceptor cell retinol dehydrogenase cause childhood-onset severe retinal dystrophy. Nature Genetics 36, 850–854 (2004).

52. Bareil, C. et al. Segregation of a mutation in CNGB1 encoding the β-subunit of the rod cGMP-gated channel in a family with autosomal recessive retinitis pigmentosa. Human Genetics 108, 328–334 (2001).

53. Pierce, E.A. et al. Mutations in a gene encoding a new oxygen-regulated photoreceptor protein cause dominant retinitis pigmentosa. Nature Genetics 22, 248–254 (1999).

54. Sullivan, L.S. et al. Mutations in a novel retina-specific gene cause autosomal dominant retinitis pigmentosa. Nature Genetics 22, 255–259 (1999).

55. Kohl, S. et al. Mutations in the Cone Photoreceptor G-Protein Subunit Gene *GNAT2* in Patients with Achromatopsia. The American Journal of Human Genetics 71, 422–425 (2002).

56. Rosenberg, T. et al. Variant Phenotypes of Incomplete Achromatopsia in Two Cousins with GNAT2 Gene Mutations. Investigative Ophthalmology & Visual Science 45, 4256–4262 (2004).

57. Thiadens, A.A.H.J. et al. Homozygosity Mapping Reveals *PDE6C* Mutations in Patients with Early-Onset Cone Photoreceptor Disorders. The American Journal of Human Genetics 85, 240–247 (2009).

58. Kohl, S. et al. A Nonsense Mutation in *PDE6H* Causes Autosomal-Recessive Incomplete Achromatopsia. The American Journal of Human Genetics 91, 527–532 (2012).

59. Piri, N. et al. A substitution of G to C in the cone cGMP-phosphodiesterase γ subunit gene found in a distinctive form of cone dystrophy. Ophthalmology 112, 159–166 (2005).

60. Kaufman, M.L. et al. Transcriptional profiling of murine retinas undergoing semi-synchronous cone photoreceptor differentiation. Developmental Biology 453, 155–167 (2019).

61. Irie, S. et al. Rax Homeoprotein Regulates Photoreceptor Cell Maturation and Survival in Association with Crx in the Postnatal Mouse Retina. Molecular and Cellular Biology 35, 2583–2596 (2015).

62. de Melo, J., Peng, G.-H., Chen, S. & Blackshaw, S. The Spalt family transcription factor Sall3 regulates the development of cone photoreceptors and retinal horizontal interneurons. Development 138, 2325–2336 (2011).

63. Onishi, A. et al. Pias3-Dependent SUMOylation Directs Rod Photoreceptor Development. Neuron 61, 234–246 (2009).

64. Aavani, T., Tachibana, N., Wallace, V., Biernaskie, J. & Schuurmans, C. Temporal profiling of photoreceptor lineage gene expression during murine retinal development. Gene Expression Patterns 23**-** 24, 32-44 (2017).

65. Tran, N.M. & Chen, S. Mechanisms of blindness: Animal models provide insight into distinct CRX-associated retinopathies. Developmental Dynamics 243, 1153–1166 (2014).

66. Roger, J.E. et al. OTX2 loss causes rod differentiation defect in CRX-associated congenital blindness. The Journal of Clinical Investigation 124, 631–643 (2014).

67. Baird-Titus, J.M. et al. The solution structure of the native K50 Bicoid homeodomain bound to the consensus TAATCC DNA-binding site.

68. Chaney, B.A., Clark-Baldwin K Fau - Dave, V., Dave V Fau - Ma, J., Ma J Fau - Rance, M. & Rance, M. Solution structure of the K50 class homeodomain PITX2 bound to DNA and implications for mutations that cause Rieger syndrome.

69. Chu, S.W., et al. Exploring the DNA-recognition potential of homeodomains. (2012).

70. Kimura, A. et al. Both PCE-1/RX and OTX/CRX Interactions Are Necessary for Photoreceptor-specific Gene Expression *. Journal of Biological Chemistry 275, 1152–1160 (2000).

71. Hughes, A.E.O., Myers, C.A. & Corbo, J.C. A massively parallel reporter assay reveals context-dependent activity of homeodomain binding sites in vivo. Genome Res 28, 1520–1531 (2018).

72. White, Michael A. et al. A Simple Grammar Defines Activating and Repressing *cis*-Regulatory Elements in Photoreceptors. Cell Reports 17, 1247–1254 (2016).

73. Gillinder, K.R. et al. Promiscuous DNA-binding of a mutant zinc finger protein corrupts the transcriptome and diminishes cell viability.

74. Wang, S. & Cepko, C.L. Photoreceptor Fate Determination in the Vertebrate Retina. Investigative ophthalmology & visual science 57, ORSFe1–ORSFe6 (2016).

75. Brzezinski, J.A. & Reh, T.A. Photoreceptor cell fate specification in vertebrates. Development 142, 3263–73 (2015).

76. Sapkota, D. et al. Onecut1 and Onecut2 redundantly regulate early retinal cell fates during development.

77. Mori, M., Ghyselinck Nb Fau - Chambon, P., Chambon P Fau - Mark, M. & Mark, M. Systematic immunolocalization of retinoid receptors in developing and adult mouse eyes. (2001).

78. Roberts, M.R., Hendrickson A Fau - McGuire, C.R., McGuire Cr Fau - Reh, T.A. & Reh, T.A. Retinoid X receptor (gamma) is necessary to establish the S-opsin gradient in cone photoreceptors of the developing mouse retina.

79. Carter-Dawson, L.D. & LaVail, M.M. Rods and cones in the mouse retina. I. Structural analysis using light and electron microscopy. J Comp Neurol 188, 245–62 (1979).

80. Vincent, A. et al. *OTX2* mutations cause autosomal dominant pattern dystrophy of the retinal pigment epithelium. Journal of Medical Genetics 51, 797 (2014).

81. Ragge, N.K. et al. Heterozygous Mutations of *OTX2* Cause Severe Ocular Malformations. The American Journal of Human Genetics 76, 1008–1022 (2005).

82. Perveen, R. et al. Phenotypic Variability and Asymmetry of Rieger Syndrome Associated with PITX2 Mutations. Investigative Ophthalmology & Visual Science 41, 2456–2460 (2000).

83. Rister, J. et al. Single-base pair differences in a shared motif determine differential Rhodopsin expression. Science 350, 1258–61 (2015).

84. Tucker, S.C. & Wisdom, R. Site-specific Heterodimerization by Paired Class Homeodomain Proteins Mediates Selective Transcriptional Responses*. Journal of Biological Chemistry 274, 32325–32332 (1999).

85. Chirco, K.R., Chew, S., Moore, A.T., Duncan, J.L. & Lamba, D.A. Allele-specific gene editing to rescue dominant CRX-associated LCA7 phenotypes in a retinal organoid model. Stem Cell Reports 16, 2690–2702 (2021).

86. Yang, H., Wang, H. & Jaenisch, R. Generating genetically modified mice using CRISPR/Cas-mediated genome engineering. Nature Protocols 9, 1956–1968 (2014).

87. Cho, A., Haruyama, N. & Kulkarni, A.B. Generation of transgenic mice. Curr Protoc Cell Biol Chapter 19, Unit 19.11 (2009).

88. Chen, S. et al. Interference of Crx-dependent transcription by ataxin-7 involves interaction between the glutamine regions and requires the ataxin-7 carboxy-terminal region for nuclear localization. Human Molecular Genetics 13, 53–67 (2004).

89. Schmidt, D. et al. ChIP-seq: using high-throughput sequencing to discover protein-DNA interactions. Methods 48, 240–8 (2009).

90. Lee, J., Myers, C.A., Williams, N., Abdelaziz, M. & Corbo, J.C. Quantitative fine-tuning of photoreceptor cis-regulatory elements through affinity modulation of transcription factor binding sites. Gene Therapy 17, 1390–1399 (2010).

91. Roy, B., Zuo, Z. & Stormo, G.D. Quantitative specificity of STAT1 and several variants. Nucleic acids research 45, 8199–8207 (2017).

92. Felix Krueger, F.J., Phil Ewels, Ebrahim Afyounian, Michael Weinstein, Benjamin Schuster-Boeckler, Gert Hulselmans, sclamons. FelixKrueger/TrimGalore: v0.6.9 - fix declaration bug (0.6.9). Zenodo (2023).

93. FastQC. (2015).

94. Langmead, B. & Salzberg, S.L. Fast gapped-read alignment with Bowtie 2. Nature Methods 9, 357–359 (2012).

95. Li, H. et al. The Sequence Alignment/Map format and SAMtools. Bioinformatics 25, 2078–2079 (2009).

96. Picard toolkit. Broad Institute, GitHub repository (2019).

97. Quinlan, A.R. & Hall, I.M. BEDTools: a flexible suite of utilities for comparing genomic features. Bioinformatics 26, 841–842 (2010).

98. Ramírez, F. et al. deepTools2: a next generation web server for deep-sequencing data analysis. Nucleic Acids Research 44, W160–W165 (2016).

99. Robinson, J.T., et al. Integrative genomics viewer. Nature Biotechnology 29, 24–26 (2011).

100. Zhang, Y. et al. Model-based Analysis of ChIP-Seq (MACS). Genome Biology 9, R137 (2008).

101. Li, Q., Brown, J.B., Huang, H. & Bickel, P.J. Measuring reproducibility of high-throughput experiments. The Annals of Applied Statistics 5, 1752–1779, 28 (2011).

102. Stark, R. & Brown, G. DiffBind: Differential binding analysis of ChIP-Seq peak data. (2012).

103. Ross-Innes, C.S. et al. Differential oestrogen receptor binding is associated with clinical outcome in breast cancer. Nature 481, 389–393 (2012).

104. Love, M.I., Huber, W. & Anders, S. Moderated estimation of fold change and dispersion for RNA-seq data with DESeq2. Genome Biology 15, 550 (2014).

105. McLean, C.Y. et al. GREAT improves functional interpretation of cis-regulatory regions. Nat Biotechnol 28, 495–501 (2010).

106. Gu, Z. & Hübschmann, D. rGREAT: an R/bioconductor package for functional enrichment on genomic regions. Bioinformatics 39(2022).

107. Müllner, D. fastcluster: Fast Hierarchical, Agglomerative Clustering Routines for R and Python. Journal of Statistical Software 53, 1–18 (2013).

108. Pagès, H. BSgenome: Software infrastructure for efficient representation of full genomes and their SNPs. (2020).

109. Bailey, T.L., Johnson, J., Grant, C.E. & Noble, W.S. The MEME Suite. Nucleic Acids Research 43, W39–W49 (2015).

110. Bray, N.L., Pimentel, H., Melsted, P. & Pachter, L. Near-optimal probabilistic RNA-seq quantification. Nature Biotechnology 34, 525–527 (2016).

111. Soneson, C., Love, M.I. & Robinson, M.D. Differential analyses for RNA-seq: transcript-level estimates improve gene-level inferences. F1000Res 4, 1521 (2015).

112. Sherman, B.T. et al. DAVID: a web server for functional enrichment analysis and functional annotation of gene lists (2021 update). Nucleic Acids Res 50, W216–21 (2022).

113. Yu, G., Wang, L.G., Han, Y. & He, Q.Y. clusterProfiler: an R package for comparing biological themes among gene clusters. Omics 16, 284–7 (2012).

114. Wu, T. et al. clusterProfiler 4.0: A universal enrichment tool for interpreting omics data. Innovation (Camb) 2, 100141 (2021).

115. Carlson, M. org.Mm.eg.db: Genome wide annotation for Mouse. (2019).

116. Virtanen, P. et al. SciPy 1.0: fundamental algorithms for scientific computing in Python. Nature Methods 17, 261–272 (2020).

117. Terpilowski, M.A. scikit-posthocs: Pairwise multiple comparison tests in Python. Journal of Open Source Software 4, 1169 (2019).

118. Goujon, M. et al. A new bioinformatics analysis tools framework at EMBL–EBI. Nucleic Acids Research 38, W695–W699 (2010).

119. Sievers, F. et al. Fast, scalable generation of high-quality protein multiple sequence alignments using Clustal Omega. Molecular Systems Biology 7, 539 (2011).

120. Waterhouse, A.M., Procter, J.B., Martin, D.M.A., Clamp, M. & Barton, G.J. Jalview Version 2—a multiple sequence alignment editor and analysis workbench. Bioinformatics 25, 1189-1191 (2009).

121. Tareen, A. & Kinney, J.B. Logomaker: beautiful sequence logos in Python. Bioinformatics 36, 2272–2274 (2019).

122. Hunter, J.D. Matplotlib: A 2D Graphics Environment. Computing in Science & Engineering 9, 90–95 (2007).

123. Jeff Reback, j., Wes McKinney, Joris Van den Bossche, Tom Augspurger, Matthew Roeschke, Simon Hawkins, Phillip Cloud, gfyoung, Sinhrks, Patrick Hoefler, Adam Klein, Terji Petersen, Jeff Tratner, Chang She, William Ayd, Shahar Naveh, JHM Darbyshire, Marc Garcia, Richard Shadrach, Jeremy Schendel, Andy Hayden, Daniel Saxton, Marco Edward Gorelli, Fangchen Li, Matthew Zeitlin, Vytautas Jancauskas, Ali McMaster, Torsten Wörtwein, Pietro Battiston. pandas-dev/pandas: Pandas 1.4.2. (2022).

124. Van Rossum, G.a.D.J., Fred L. Python reference manual, (Centrum voor Wiskunde en Informatica Amsterdam, 1995).

125. Waskom, M.L. seaborn: statistical data visualization. Journal of Open Source Software 6, 3021 (2021).

